# Interbacterial AI-2 communication drives stage-specific genetic programs to support *Salmonella* colonisation in the murine gut

**DOI:** 10.1101/2025.07.28.667109

**Authors:** Anmol Singh, Abhilash Vijay Nair, R. S. Rajmani, Dipshikha Chakravortty

## Abstract

The cooperative response through AI-2 allows *Salmonella* to coordinate and regulate its community behaviours crucial for its survival and infection. Understanding these mechanisms is essential for developing strategies to interfere with *Salmonella’s* ability to cause disease. In this study, we report that *Salmonella* Typhimurium uses AI-2 signaling to modulate the expression of chemotaxis and motility-associated genes and the formation of flagella on the bacterial cell surface. The AI-2 acts as a chemoattractant, and the cascade mediates the chemotactic response of *Salmonella.* Such regulations assist in the attachment and invasion of *Salmonella* into intestinal epithelial cells both *in vitro* and *in a* murine model. Furthermore, we determined that HilD, known to regulate motility by induction of a chemoreceptor and SPI-1 in *Salmonella*, along with key SPI-1 genes, is underexpressed in the absence of LuxS/AI-2 signaling, which unravels a complex regulatory network by AI-2 in *Salmonella*. Beyond the initial phase, this signaling facilitates survival in the presence of polymyxin B and supports intracellular survival and persistence within epithelial cells by regulating the *pmrD/AB* system, thereby evading the antimicrobial peptide immune response. Additionally, our findings show that *Salmonella* activates its LuxS/AI-2 signalling cascade within the Caco-2 epithelial cells. Complimentary observation through transcriptomic profiling revealed that LuxS/AI-2 signalling governs the coordinated regulation of genes implicated in distinct phases of *Salmonella* pathogenesis. Lastly using a mouse model, we show that AI-2 signaling is critical for organ colonization and virulence of *Salmonella,* and inhibition of LuxS/AI-2 signaling in combination with antibiotics can be an alternative therapeutic approach. Thus, our study dissects an integrated regulatory mechanism by which AI-2 signaling mediates critical processes during multiple stages of *Salmonella* Typhimurium pathogenesis.

## Introduction

Bacteria were initially thought to be solitary microbes that led an independent life cycle of multiplying and increasing their numbers, and some of them could cause life-threatening infections. However, a series of discoveries led to unravelling the mode of communication in these little beings referred to as quorum sensing (1–3). This process involves the production and secretion of chemical signals that are sensed by the neighbouring cells in the bacterial community and orchestrate a myriad of genes and pathways, resulting in coordinated behaviours such as adaptation, virulence and other critical community behaviours underscoring the advantages of communal living among bacterial populations (4, 5). Bacterial pathogenesis is very well governed by this communication paradigm. The pathogen, upon entry into the host, has to decide to stay and colonise, or utilise the host nutrients and run or employ all its virulence strategies without worrying about the repercussions. Interestingly, the community communicates, coordinates and makes the decision on the microbial pathogenesis (6, 7). Bacterial pathogens are the sole cause of serious bacterial infections (SBIs) worldwide, with an estimated 7.7 million deaths globally, with a rise in antimicrobial resistance, which further worsens the situation(8). *Salmonella* enterica serovar Typhimurium (STM) is a prominent etiological agent of foodborne illnesses and a leading contributor to non-typhoidal salmonellosis (NTS)(9). It faces very long odds while seeking to colonise and breach the intestinal barrier, including acidic pH, limited nutrient availability, bile salt toxicity, and competition from gut microbiota. It is critical for the bacteria to establish and increase in number, reaching the infectious dose to invade the host epithelial cells. Upon breaching the epithelial barrier, it is engulfed by macrophages in the lamina propria, facilitating its systemic dissemination throughout the host (10–13). With the fluctuations of extreme conditions, *Salmonella* needs to be adept at sensing, responding, and adapting to unprecedented situations.

*Salmonella* uses the autoinducer-2 (AI-2) chemical messenger for their communication. This intercellular communication, called quorum sensing, enables *Salmonella* to regulate and synchronize their behaviors when present at high cell densities(14–16). The AI-2 in *S*. Typhimurium is a byproduct of the activated methyl cycle (AMC), where initially, the S-adenosyl methionine(SAM) is converted into S-adenosylhomocysteine(SAH) by methyl transferases (17). The SAH is subsequently converted to the precursor of AI-2 molecule 4,5 dihydroxy-2,3-pentanedione (DPD) by the enzymes Pfs and LuxS, respectively(18). Finally, DPD molecules autocyclize into (2R,4S)-2-methyl-2,3,3,4-tetrahydroxytetra-hydrofuran (AI-2) molecules(19, 20).

Although the LuxS/AI-2 quorum-sensing signaling pathway is well characterized, its physiological significance remains only partially understood in *Salmonella* pathogenesis despite being a thoroughly established system. Studies suggest that *Salmonella* quorum sensing regulates virulence, biofilm formation, motility, and antimicrobial resistance by regulating *Salmonella* Pathogenicity Island-1, flagella, and antioxidative genes(21–23). Further, there is evidence of involvement of AI-2 in motility and chemotaxis in the Ag43-dependent aggregation of *E. coli* and its roles in *E. coli* gut colonization(24). The study also shows that chemotaxis towards self-produced AI-2 provides *E. coli* a fitness advantage during gut colonization, hinges on fructoselysine metabolism(25). AI-2, a molecule produced and detected by numerous bacterial species, has also been found to be mimicked and synthesized by eukaryotic cells(26). Research has demonstrated that several bacterial species exhibit a chemotactic response to AI-2, highlighting its crucial role in forming intricate multispecies communities(27). Notably, AI-2 is thought to influence the bacterial community structure in the mammalian gut following antibiotic-induced dysbiosis and to bolster colonization resistance against specific enteric pathogens(28). Thus, it is apparent that *Salmonella* communication is a key to its pathogenesis. Despite these findings, significant gaps remain in understanding the advantages of communal living and how AI-2 signaling benefits *Salmonella* in physiologically relevant conditions.

In our present study, we elucidate that the LuxS/AI-2 signaling cascade governs the processes of the initial attachment and invasion into intestinal epithelial cells *in vivo* in mice and *in vitro* by regulating chemotaxis and flagellar motility. Furthermore, LuxS/AI-2 signaling assists in intracellular survival by regulating the *pmrABC* operon, which provides resistance against antimicrobial peptides. As an alternative approach, inhibition of AI-2 signaling in combination with antibiotics reduced *Salmonella* organ burden and delayed mortality in mice models. We investigated the underlying mechanism of bacterial communal living and communication, driving key stages of *Salmonella* pathogenesis within the host.

## Materials and Methods

### Bacterial strains and Growth conditions

The wild-type *Salmonella* enterica serovar Typhimurium strain 14028S (STM WT), generously provided by Professor Michael Hensel from the Max Von Pettenkofer-Institute for Hygiene and Medical Microbiology in Germany, was employed in all experiments of this study. Bacterial strains were revived on LB agar, with or without antibiotics, and LB broth cultures of wild-type, knockout, and complemented strains were incubated at 37°C (170 rpm) using an orbital shaker. Wherever necessary, antibiotics were used, including Kanamycin (50 μg/ml), Chloramphenicol (25 μg/ml), and Ampicillin (50 μg/ml).

### Bacterial gene knockout and strain generation

*Salmonella enterica* serovar Typhimurium strain 14028 and isogenic mutants were used for all experiments. *luxS*, *lsrB*, *lsrK,* and *lsrR* gene knockout and *luxS* complement strains were made in our previous study **(S Table 1)**. Briefly, all the knockout generation was done as described previously in one gene step chromosomal gene inactivation method demonstrated by Datsenko and Wanner (29).

Briefly, for the complement strain, the double-digested PCR amplified purified *luxS* gene (containing BamH1 and HindIII restriction site sequences) was used to clone in double-digested pQE-60. The respective ligated vector was transformed into the STM Δ*luxS* strains to generate the complement strains. For the reporter plasmid *plsr*-GFP generation, 256 bp of upstream of *lsrA* were cloned into the promoter less vector pUA66. Finally, the reporter plasmid was transformed into the STM WT and STM Δ*luxS*.

### Cell culture and maintenance

Caco-2 epithelial cell lines were maintained in Dulbecco’s Modified Eagle Medium (DMEM, Lonza), supplemented with 10% FBS (Gibco), 1% non-essential amino acids, 1% sodium pyruvate, and 1% penicillin-streptomycin (Sigma-Aldrich) in a humidified incubator with 5% CO2 at 37 °C. The cells were seeded into the respective cell culture plates for each experiment to conduct intracellular survival assays and intracellular gene expression experiments.

### Gentamicin protection assay

Caco-2 cells were seeded in tissue culture plates for infection. Cells were infected with STM WT, STM Δ*luxS,* STM Δ*lsrB,* STM Δ*lsrK,* STM Δ*lsrR,* and *STM*Δ*luxS:luxS* (log phase culture growing overnight in LB broth). The multiplicity of Infection (MOI) 20 was used for the intracellular survival assay and *Salmonella* gene expression study (qRT-PCR). Following the infection of STM strains into the cell line, the plate was spun for 5 minutes at 600 rpm using a Rota-Superspin R–NV swing bucket centrifuge. After that, the plate was incubated for 25 minutes at 37 °C with 5% CO_2_ in a humidified incubator. Then, the infected cells were washed with 1X PBS twice and treated with a 100 μg/ mL concentration of gentamycin. After 1 hour, the media were changed with a reduced concentration of gentamycin (25 μg/ mL) and incubated until time point. The time points 2h, 6h, and 16h were taken for the qRT-PCR sample and 2h and 16h for the intracellular survival assay.

DPD supplementation (of 25 μM, 50 μM,100 μM, and 200μM DPD) or spent media of STM WT collected at 3h in LB growing treated STM *ΔluxS,* and LuxS/AI-2 signaling inhibition(30, 31), the inhibitor (Z)-4-bromo-5-(bromomethyllene)-3-methylfran-2(5H), known as BF-8 treatment, was given to the STM WT culture, which was used for the gentamycin protection assay to study the adhesion, invasion, and intracellular proliferation of STM.

### Invasion and intracellular survival assay

The cells were lysed with 0.1% triton-X 100 in PBS at specific time points post-infection. For the invasion, CFU was determined at 2h post-infection samples and CFU in pre-inoculum. Percent invasion was calculated by using the formula- [CFU at 2 hours]/ [CFU of pre-inoculum] ×100. For the fold proliferation, CFU was determined at 2h and 16h post-infection samples. Fold proliferation was determined using the formula [CFU at 16 hours]/ [CFU at 2 hours].

### Adhesion assay

Adhesion assay was done as gentamycin protection assay with modification. Infections were given to Caco-2 cells at MOI 20 and incubated for 10 minutes at 37 °C in a humidified incubator with 5% CO2. Thereafter, washing with 1X PBS twice to eliminate any loosely adherent bacteria. After lysing the mammalian cells with 0.1% Triton-X 100, samples were taken to release the bacteria into the sample. After the collected samples were plated on LB agar plates at the appropriate dilutions and incubated at 37 °C, the CFUs were enumerated. The following formula was used to calculate the percentage adhesion = [(CFU at 10 min post-infection)/(CFU of the pre-inoculum)]× 100.

### *In vitro* motility assay

2 µl of bacterial culture was spotted onto the agar plates supplemented with 1% casein enzyme hydrolysate, 0.5% yeast extract, 0.5% NaCl, and 0.5% glucose with 0.3 % agar (swim agar plates) and 0.5% agar plates (swarm agar plates). The swim agar plates were incubated at 37°C for 6h, and swarm agar plates were incubated for 12h. The diameters of the motility halos were measured at least three angles of three replicate plates, and then images were taken using a BioRad chemidoc.

### Chemotaxis assay

Chemotaxis assay was done by using µ-Slide Chemotaxis, ibidi. µ-Slide consists of two distinct opposing liquid chambers or reservoirs separated by a small liquid transition zone of 1 mm. Overnight grown STM WT, STM *ΔluxS*, and STM *ΔlsrB* were followed by two washes with motility buffer (MB; 10 mM KPO4, 0.1 mM EDTA, and 67 mM NaCl, pH 7), and then resuspension to a final OD600 to 0.3 in MB supplemented with 3.5% glucose (MBg)(24). To reduce metabolic activity, cells were chilled for 20 minutes. A motility buffer was applied to the transition zone. The first reservoir was completely filled with the bacterial cell suspension, the second reservoir with STM WT spent media (Collected at 3h of LB growth), synthetic DPD molecule, and the known chemoattractant L-serine. In a different experiment, the numbers of bacteria that transmigrated from the first reservoir into the second reservoir were quantified by CFU analysis and bright field microscopy (Leica microscope SP8 falcon, 63X).

### RNA isolation and RT-qPCR

For the gene expression study in Luria Bertani (LB) media, overnight-grown culture in LB media was subjected to a subculture (at 1:100) in LB. At 3h, 6h,9h, and 12h, bacterial cell pellets were collected in TRIzol (from TaKaRa, RNA isoPlus-9109) and kept at -80°C. Total RNA was isolated by chloroform extraction followed by Isopropanol precipitation. To evaluate the quality, the amount of RNA was quantified in nano drops and examined on a 2% agarose gel. To produce cDNA, 2 μg of RNA sample was treated with DNase I at 37°C for 1 hour (TaKaRa), followed by heating for 10 minutes at 65°C. Then, DNA-free RNA samples were used to make cDNA by using the PrimeScript RT reagent Kit provided by TaKaRa (Cat# RR037A).

The gene expression of *Salmonella* upon Infection in Caco-2 infection was given at MOI 20. At 2h, 6h, and 16h post-infection, samples were collected in TRIzol (from TaKaRa, RNA isoPlus-9109) and kept at -80°C. Total RNA isolation and cDNA were synthesized using a similar protocol described in the kit provided, TRIzol (from TaKaRa, RNA isoPlus-9109) and TaKaRa (Cat# RR037A) reagents according to the manufacturer’s protocol. The list of expressions Primer is provided in **S Table 2**. Quantitative PCR was carried out using a SYBR Green Q-PCR kit (Takara), and the ΔΔCt method was used for quantification.

### RNA sequencing and analysis

Bacteria were grown till mid-log phase and RNA extraction was performed using Qiagen RNeasy Mini kit (Cat. No. 74104) with a few modifications. The RNA samples were quantified on Nanodrop and Qubit flurometer using appropriate standards. Appropriate dilutions were loaded on a High sensitivity RNA screen tape and run on Agilent Tape station 4200 to determine the RNA integrity number (RIN). For Library preparation, 1000ng of RNA sample was used for ribodepletion by using NEB Next rRNA depletion kit Bacteria (Cat. no. NEB E7850X), which includes Probe hybridization of RNA, RNase H digestion, DNase 1 digestion according to the manufacturer’s protocol. The NEBNext® Ultra II Directional RNA Library prep kit for Illumina (Cat. no. E7760L), was used to construct double-stranded cDNA libraries from the rRNA depleted RNA, using RNA fragmentation, first strand cDNA synthesis with random primers, second strand cDNA synthesis, end repair of double stranded cDNA, adapter ligation, removal of excess adapter using sample purification beads, PCR enrichment of adapter ligated DNA and clean-up of PCR products as instructed by the manufacturer. The cleaned libraries were quantified on Qubit®flurometer and appropriate dilutions loaded on a D1000 screen tape to determine the size range of the fragments and the average library size.

The original raw data from the Illumina platform is transformed into sequenced reads by base calling. Raw data are recorded in a FASTQ file containing sequence information (reads) and corresponding sequencing quality information. The quality of the reads was assessed using Fast QC v 0.11.9 before proceeding with the downstream analysis(32). The reads were passed through SortMeRNAv4.3.7 (33) and were used against all eight rRNA databases provided to filter and remove rRNA reads. The trimmed reads were then passed through fastp (34) to remove adaptors, low-quality bases (Q< 36 bp). For mapping and quantification, *Salmonella* Typhimurium 14028s reference genome and annotation were downloaded from (https://ftp.ncbi.nlm.nih.gov/genomes/all/GCF/000/022/165/GCF_000022165.1_ASM2216v1/). HISAT2 (35) was used to create the reference index, and the trimmed & filtered reads were mapped to the indexed genome using HISAT2. Quantification was performed with feature Count (36). A total of 4728 features were assigned in the raw count file. The raw count file is made into raw counts for individual sets, and the rest of the differential analysis is done according to the sets. The library sizes are plotted for individual sets. The transcripts with low read counts (< count of at least 10 for a minimum of 2 samples) were removed. The filtered read counts were normalized by transforming to the rlog transformation function (rlog) of DESeq2 (37) and are used for further analysis. Differential gene expression was analysed using the DESeq function from the DESeq2 package with a false discovery rate (FDR) cut-off of ≤ 0.05 and a minimum expression log2-fold change (FC) of ≥ 2/1. The counts are illustrated with a volcano plot. The top genes with the largest log2fold change are plotted as a heatmap.

### Transmission electron microscopy

Flagella were visualized using the protocol described previously(38, 39). Briefly, overnight bacterial cultures were subcultured in LB media and incubated until it reached an OD of 0.1. During the subculture, STM WT was treated with AI-2 signaling inhibitor BF-8, and STM *ΔluxS* with STM WT spent media and synthetic DPD. The bacterial cultures were centrifuged at 2000 rpm for 10 min at 4◦C. The bacterial cells were washed with 1XPBS twice, and finally, the cells were resuspended in 100 µl of 1X PBS. 10 µl of the cell suspension was added on a copper grid, air dried, stained with 1% uranyl acetate for 30 s, and visualized under a transmission electron microscope (Thermo-Fischer Scientific).

### Immunofluorescence

After appropriate post-infection incubation time in the adhesion assay, the media was removed, and the cells were washed with 1X PBS and fixed with 3.5% Paraformaldehyde for 10 min. The cells were then washed with 1X PBS, incubated with the required rabit-raised anti-*Salmonella* primary antibody in a buffer containing 2% BSA, and incubated for 1 hour at room temperature or overnight at 4℃min. After washing with 1X PBS, the secondary antibody tagged to a fluorochrome was added and incubated. For Caco-2 epithelial cells, FM464 dye was used. The coverslips were washed with PBS, mounted on a clean glass slide using mounting media containing an anti-fade reagent, and observed under the confocal microscope (Leica microscope SP8 falcon, at 63X oil immersion).

Mice intestine histochemistry samples were incubated with the required rabbit-raised anti-*Salmonella* primary antibody in a buffer containing 0.01% saponin and 2% BSA, and incubated for 1 hour at room temperature or overnight at 4℃min. After washing with 1X PBS, the secondary antibody tagged to a fluorochrome was added and incubated for 1h at room temperature. The dye DAPI was used to stain the intestinal nucleus. 40X oil immersion was used to study the mice intestine histopathology (Zeiss 710 microscope, at 63X oil immersion).

### Polymyxin treatment and survival assay

Polymixin MIC was determined by using microbroth dilution methods. Polymixin treatment (sub-MIC concentration 0.5 µg/ml) was given to the overnight STM strains (10^7^ cfu) in LB media. At 3h post-treatment, CFU was determined by spread plate methods. For the time-dependent survival assay, we collected the treated sample at 3h, 6h, 9h, and 12h and calculated the CFU. For the mRNA expression study, samples were collected during mid-log phase (6h) in Trizol and kept at -80℃ for further RNA isolation.

### Flow cytometry

Reporter strain STM WT-*plsr*-GFP and STM *ΔluxS*-*plsr*-GFP were grown in LB media, and samples were taken out at different time points and fixed with 3.5% paraformaldehyde. The positive GFP percentage population was determined in the sample by using BD FACS Verse.

### *In vivo* animal experiment

Male C57BL/6, aged 5 to 6 weeks, were obtained from the Central Animal Facility, IISc.

### Invasion assay

5–6 weeks old C57BL/6 mice were infected by orally gavaging 107 CFU of STM WT, STM *ΔluxS*, STM *ΔlsrB*, STM *ΔlsrK*, STM *ΔlsrR,* and STM *ΔluxS:luxS*. For invasion assay intestine was isolated 6 h post-infection, and CFU was enumerated on differential and selective SS agar by serial dilution followed by plating. BF-8 (10 µg/ml) treated STM WT culture and DPD supplementation (100 μM DPD) or STM WT spent media treated STM *ΔluxS* was used for invasion study in mice.

### *In vivo,* competition assay

Competition assay was done between STM WT and STM*ΔluxS*/STM*ΔlsrB*/STM*ΔlsrK*. Infection was given by oral gavage with 107 CFU/ml of culture in a 1:1 ratio. Post five days of infection, the intestine (Peyer’s patches), MLN (mesenteric lymph node), spleen, liver, and blood were aseptically extracted to examine the colonization in the organs. The CFU was enumerated on differential and selective *Salmonella*-*Shigella* (SS) agar and respective antibiotics containing LB agar plates. The competitive index was calculated for a WT strain and knockout strains by dividing the ratio between CFU (Knockout strain) and CFU (STM WT) by the ratio of both strains in the inoculum.

### *In vivo* infection and BF-8/antibiotics treatment

5–6 weeks old C57BL/6 mice were infected by orally gavaging 10^7^ CFU of STM WT. Post 3^rd^ day of infection, one cohort of mice was sacrificed for the organ burden of *Salmonella*. The rest of the cohort of mice were treated with either 2mg/kg of body weight of BF-8, ciprofloxacin, ceftazidime, or a combination of BF-8 with both antibiotics intraperitoneally for three consecutive days. After the 3rd day of treatment, mice were sacrificed the next day, and the organ burden was determined by CFU enumeration.

### Mice survival assay

The mice were given 10^7^CFU of STM WT (overnight grown cultures) orally to compare the survival and weight alterations of mice post-infection and BF-8 /antibiotics treatment. Post 3^rd^ day of infection, mice were treated with either 2mg/kg of mouse weight of BF-8, ciprofloxacin, ceftazidime, or a combination of BF-8 with both antibiotics, intraperitoneally for three consecutive days. The mice were observed every day to determine their survival and weight, and the results were expressed as a percentage of survival (Kaplan-Meier curve) and weight reduction.

All experiments comply with the rules set forth by the Indian Institute of Science, Bangalore’s IEAC. The approved protocol number is CAF/ Ethics/852/2021. The Institutional Animal Ethics Committee approved every animal experiment, and the National Animal Care Guidelines were scrupulously followed.

### Statistical analysis

As mentioned in figure legends, each experiment has been independently repeated two to five times. GraphPad Prism 8.4.3 was utilized for all statistical analyses. As the figure legends state, the statistical analyses included an unpaired, two-tailed Student’s t-test, One-way ANOVA with Dunnet’s post hoc test, Two-way ANOVA with Tukey’s post hoc test. A non-parametric one-way ANOVA (Kruskal Wallis) test with Dunn’s post-hoc test was performed for the animal experiment. P-values less than 0.05 were regarded as significant. The analysis is presented as mean ±SD or mean ±SEM with information on group sizes and p values mentioned in the respective figure legends.

## Results

### Autoinducer-2 signaling governs bacterial motility and flagellar organization by modulating chemotaxis and flagella-related genes

We first investigated whether *Salmonella* LuxS/AI-2 signaling is involved in motility. As reported, the aggregation of *E. coli* towards AI-2 depends on motility and chemotaxis(24),we have also observed that upon deletion of *luxS*, *lsrB*, and *lsrK*, swim and swarm motility were hampered compared to STM WT **(Fig.1A, B; S1A, B)**.

**Fig. 1.**
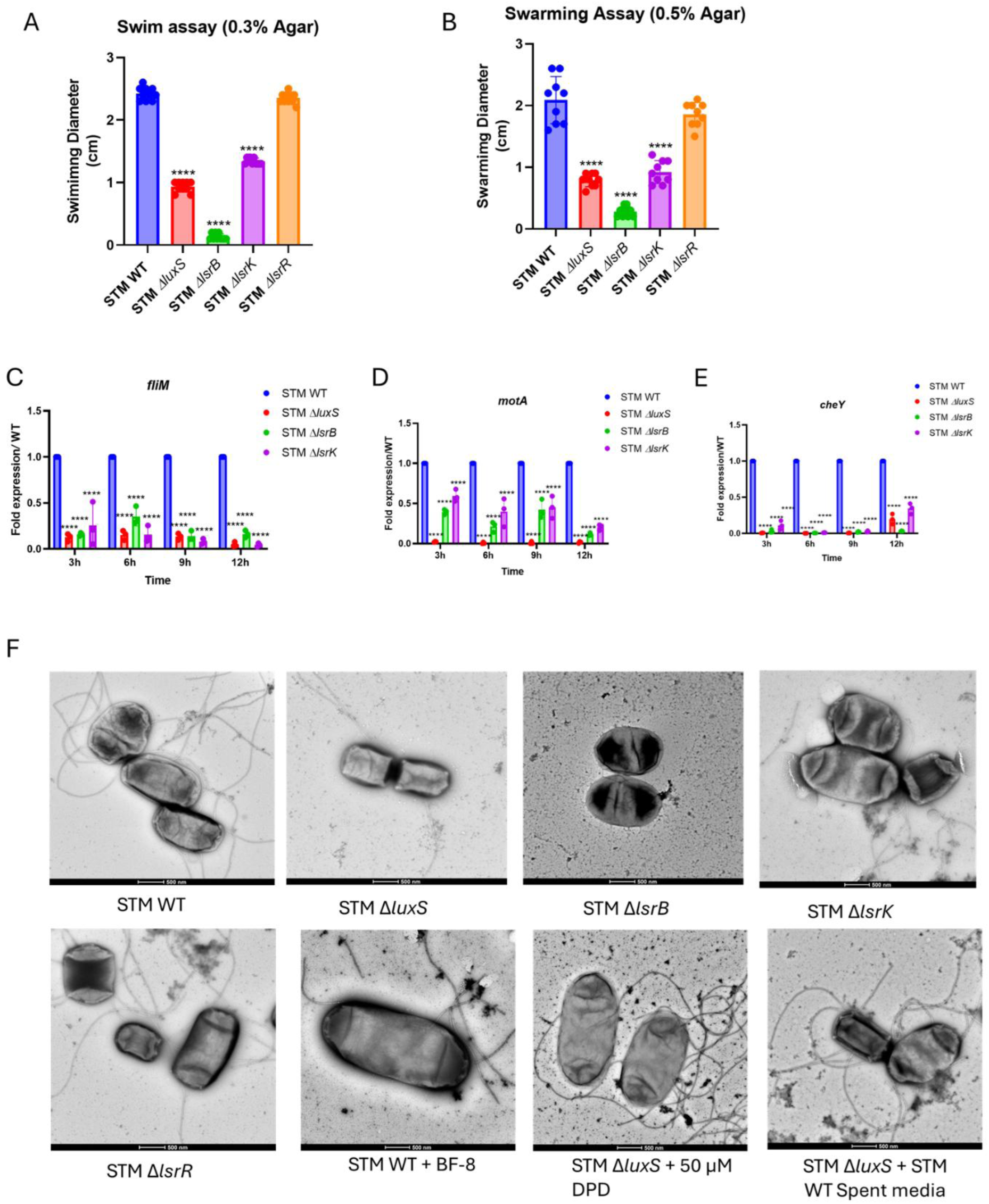
Autoinducer-2 signaling governs bacterial motility and flagellar organization by modulating chemotaxis and flagella-related genes. **(A)** Swim motility, **(B)** Swarm motility assay. Measurement has been taken at three angles per plate. Represented as Mean+/-SD of N=3, n=3. **(C)** mRNA expression of *fliM*, *motA,* and *cheY* STM strains *in vitro* growth in LB. Represented as Mean+/-SD of N=3, n=3, **(D)** Transmission electron microscope representative image of STM strains. Represented as Mean+/-SD of N=2, n>10. One-way ANOVA with Dunnet’s post-hoc test was used to analyze the data; p values **** p < 0.0001, *** p < 0.001, ** p<0.01, * p<0.05. Two-way Anova was used to analyze the grouped data; p values **** p < 0.0001, *** p < 0.001, ** p<0.01, * p<0.05

Many motile bacteria control their swimming direction by regulating the sense of flagellar rotation in response to sensory cues. FliM is a key component of the flagellar motor’s switch complex, along with FliG and FliN, which is thought to interact with one of the chemotaxis proteins, CheY(40, 41). The result of these interactions is to induce clockwise rotation of the flagellar motors and tumbling of the cell. We hypothesized that Lux/AI-2 signaling, due to its virtue of orchestrating gene expression, might regulate the expression of these genes to modulate motility. We observed that *in vitro* LB media, *fliM*, *cheY,* and *motA* (regarded as the stator of the flagellar motor)(42), mRNA expression was downregulated in STM *ΔluxS*, STM *ΔlsrB*, STM *ΔlsrK* compared to STM WT **(Fig.1C-E)**. Following this, we noted that genes involved in transcription regulation of flagella assembly *flhD, fliA, fliC, fljB*,(43) were also downregulated in the absence of AI-2 accumulation and phosphorylation **(S1 C-F)**. We further observed that in the absence of AI-2 sensing and signaling, the number of flagella is reduced on the surface of STM *ΔluxS* and STM *ΔlsrK*, while there were no flagella on the surface of *STM ΔlsrB*. We further validated with inhibition of AI-2 signaling using BF-8, a competitive inhibitor of DPD in STM WT and supplementing with exogenous DPD and STM WT spent media to STM *ΔluxS.* We observed that BF-8 treatment reduced and DPD or spent media treatment increased the flagella number (Fig.1F). From these data, we concluded that LuxS/AI-2 signaling governs motility and flagellar organisation via regulating flagella and chemotaxis-related gene expression.

### AI-2 sensing in *Salmonella* assists in chemotaxis and enhances the attachment to Caco-2 epithelial cells

Studies show that in *E.coli,* AI-2 binds LsrB, and L-serine binds Tsr chemoreceptor, and both LsrB and Tsr are necessary for sensing AI-2, but not for AI-2 uptake. This suggests that LsrB and Tsr interact directly in the periplasm within *E.coli*(27). We speculate that the LsrB receptor in *Salmonella* may act as a chemoreceptor and aid in the attraction to the AI-2 molecule. We used a µ-Slide Chemotaxis chamber (ibidi) for the chemotaxis assay. DPD, STM WT spent media, and serine were used in the second reservoir of the chamber. Serine was used as a positive control as a chemoattractant. We found that the number of STM WT transmigrated towards spent media, DPD, and serine was higher compared to STM *ΔluxS* and STM *ΔlsrB* **(Fig.2 A-C)**. The explanation for the reduced transmigration of STM *ΔluxS* and STM *ΔlsrB* toward serine is due to reduced flagella activity. These data suggest that LuxS/AI-2 signaling controls flagellar motility and chemotaxis.

**Fig. 2.**
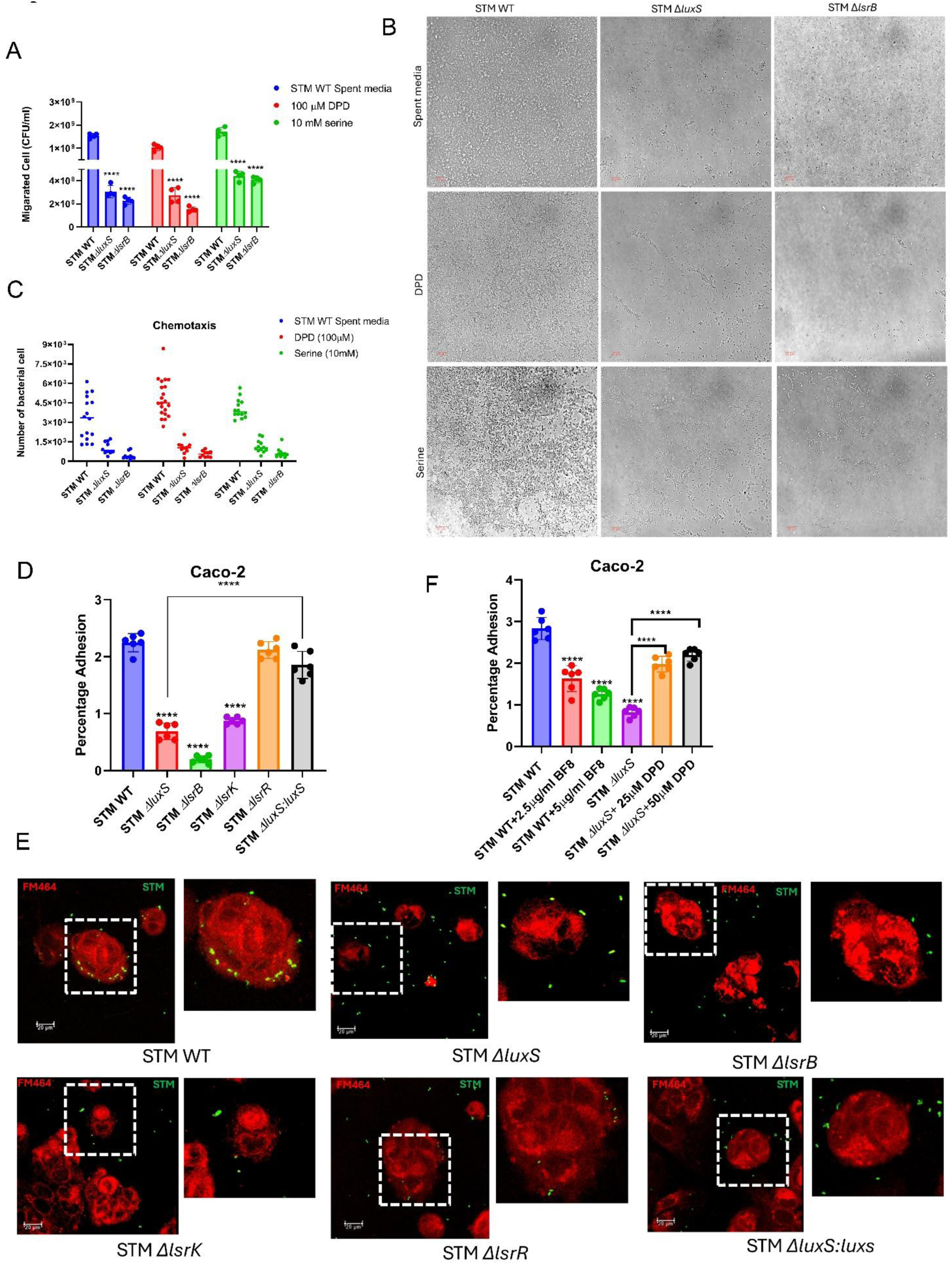
AI-2 sensing in *Salmonella* assists in chemotaxis and enhances the attachment to Caco-2 epithelial cells. **(A)** CFU analysis of transmigrated STM strains towards STM WT spent media, DPD, and known chemoattractant serine. The experiment was performed in an ibidi chemotaxis chamber. Represented as Mean+/-SD of N=3, n=2. **(B)** Bright-field microscopy images of transmigrated bacteria. **(C)** Analysis of images using ImageJ software to count the number of bacteria per image. Each dot represents per image. The image is represented as Mean+/-SD of N=3, n>10. **(D)** Adhesion assay of STM WT, STM *ΔluxS*, STM *ΔlsrB*, STM *ΔlsrK,* STM *ΔlsrR,* and STM *ΔluxS:luxS* in Caco-2 epithelial cells. Represented as Mean+/-SD of N=3, n=3. **(E)** Immunofluorescence imaging to study the adhesion to Caco-2 cells of STM WT and knockout strains, here FM464(red) is used to stain the lipids for Caco-2, and Anti-*Salmonella* (LPS) (Alexafluor-488 tagged secondary antibody used-Green) for STM. Representative image of N=3, n>10. **(F)** Adhesion assay of STM WT and STM *ΔluxS* treated with BF-8 and DPD, respectively. Represented as Mean+/-SD of N=3, n=3. One-way ANOVA with Dunnett’s post-hoc test was used to analyze the data; p values **** p < 0.0001, *** p < 0.001, ** p<0.01, * p<0.05. Two-way Anova was used to analyze the grouped data; p values **** p < 0.0001, *** p < 0.001, ** p<0.01, * p<0.05.

Replicating laboratory conditions that accurately reflect the natural microbial habitat during infection presents significant challenges. Consequently, fluid flow, a defining characteristic of natural environments, is often ignored in experimental setups. However, with the development of microfluidic technology coupled with high-resolution imaging, it is now possible to understand environmental features that affect bacterial pathogenesis(44). But this natural flow can remove the autoinducer molecule and hamper the signaling. A study revealed that *S. aureus* produces enterotoxin B in crypts, deepening them and insulating the bacteria from flow, facilitating QS-regulated pathogenicity(45). Similarly, *V. cholerae* activates QS within, but not outside, intestinal crevices, highlighting the role of topography in virulence(46, 47). *Salmonella* is a pathogen that primarily infects the ileum, the distal part of the small intestine(48). *Salmonella* has evolved mechanisms to gain access, attach, and infect the intestinal epithelial cells of the crypt to survive and replicate. We speculate that *Salmonella* is attracted towards AI-2 molecules of the intestinal crypts, where fluid flow is resisted, and it can answer why *Salmonella* moves to the distal part of the intestine. Epithelial cells release an AI-2 mimic in response to bacteria, and bacteria might benefit from this AI-2 mimic via their AI-2 quorum-sensing receptors(26). We were curious whether LuxS/AI-2 modulates the attachment to epithelial cells. Our adhesion assay suggests that STM *ΔluxS*, STM *ΔlsrB*, and *ΔlsrK* compromised their ability to adhere to Caco-2 epithelial cells **(Fig. 2D, E, S2A).** On the other hand, STM *ΔlsrR* behaves similarly to STM WT. Upon inhibition of LuxS/AI-2 signaling with BF-8 in STM WT and exogenous DPD and STM WT spent media treatment to STM *ΔluxS,* AI-2 sensing and signaling enhance the attachment to epithelial cells **(Fig. 2F).**

### *Salmonella* exploits its LuxS/AI-2 mediated communication to invade intestinal epithelial cells

Adhesion and invasion represent the crucial step of successfully establishing infection by bacterial pathogens and cellular invasion, which is followed by intracellular multiplication, dissemination to other tissues, or persistence(39, 49, 50). Having observed that LuxS/AI-2 enhances the attachment to Caco-2 cells, we also conjecture that LuxS/AI-2 signaling regulates the invasion of *Salmonella* into the epithelial cells. We infected the Caco-2 epithelial cell with STM stains and observed that STM WT, *luxS* complemented in STM *ΔluxS,* and STM *ΔlsrR* invade more efficiently compared to STM *ΔluxS*, STM *ΔlsrB*, and *ΔlsrK* **(Fig. 3A)**. Further, we treated the STM *ΔluxS* with exogenous DPD and STM WT spent media and STM WT with inhibitor BF-8 followed by infection in Caco-2 epithelial cells. Exogenous DPD and spent media treatment recovered the invasion compared to untreated STM *ΔluxS,* while upon inhibition with BF-8, the STM WT invasion was reduced **(S3 A-C)**. Next, to validate our *in vitro* cell line results, we then studied the invasion of the knockout strains into the intestine of C57BL/6 mice **(Fig. 3B).** Interestingly, STM *ΔluxS*, STM *ΔlsrB*, and *ΔlsrK* invaded significantly less into the Peyer’s patches of the C57BL/6 mice **(Fig. 3C)**.

**Fig. 3.**
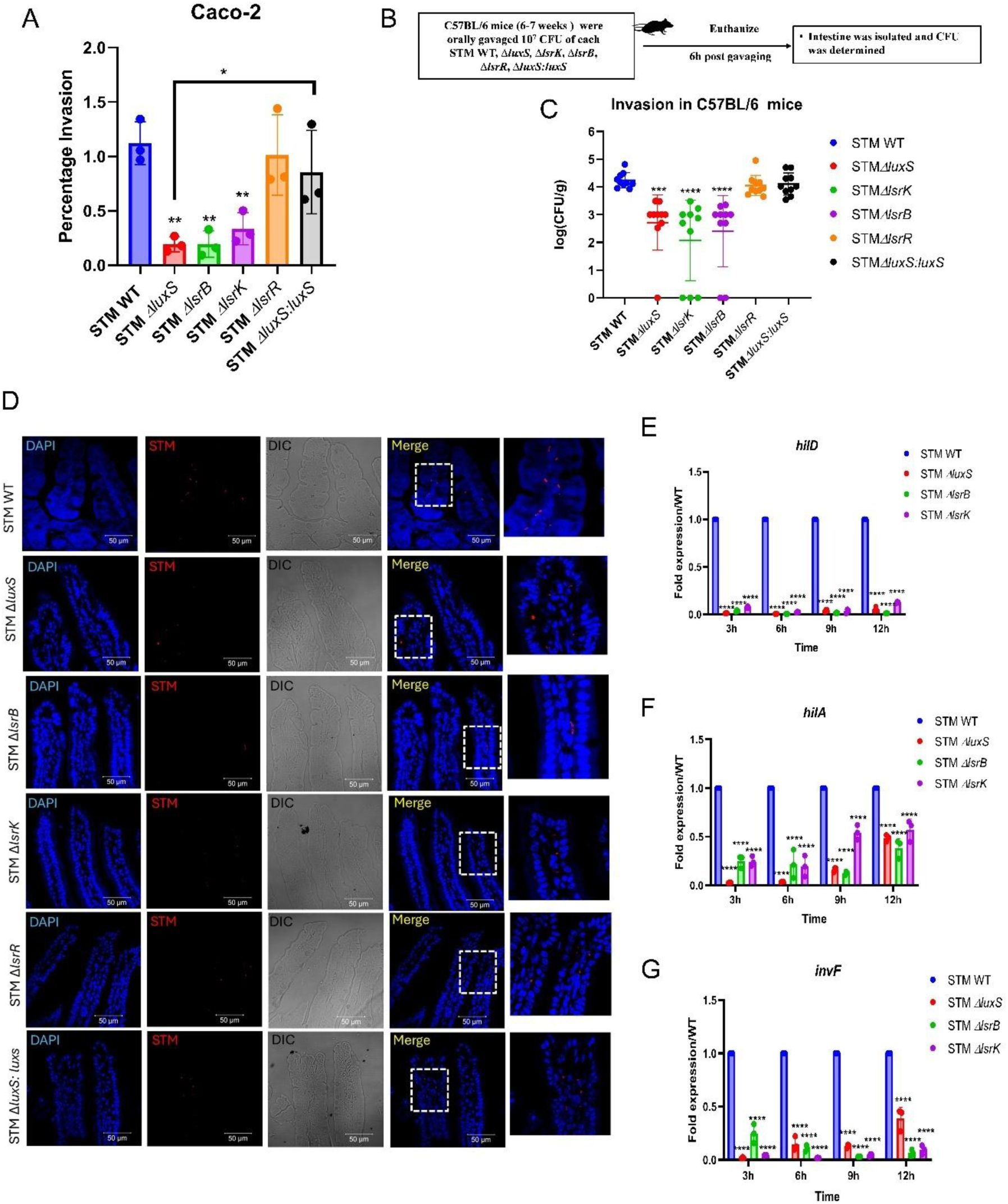
*Salmonella* exploits its LuxS/AI-2 mediated communication to invade intestinal epithelial cells. **(A)** Percentage invasion of STM WT, STM *ΔluxS*, STM *ΔlsrB*, STM *ΔlsrK,* STM *ΔlsrR,* and STM *ΔluxS:luxS* in Caco-2 epithelial cells. Represented as Mean+/-SEM of N=3, n=3. **(B)**Experimental schematic representation of the invasion assay in C57BL/6 male mice. **(C)** STM WT, STM *ΔluxS,* STM *ΔlsrB*, STM *ΔlsrK,* STM *ΔlsrR,* and STM *ΔluxS:luxS* invasion assay in mice; organ burden in the intestine post 6h of infection. N=2, n=5.**(D)** Immunofluorescence of histopathological sections of mice intestine (Peyer’s patches) to study invasion, DAPI is used to stain the nucleic acids in the cells, and Anti-*Salmonella* (LPS) (Cy3 tagged secondary antibody used-Red). mRNA expression of **(E)** *hilD,* **(F)** *hilA*, **(G)** *invF,* in STM WT, STM *ΔluxS,* STM *ΔlsrB*, and STM *ΔlsrK in vitro* LB media. Represented as Mean+/-SD of N=3, n=3. Represented as Mean+/-SD of N=2, n>10. One-way ANOVA with Dunnet’s post-hoc test was used to analyze the data; p values **** p < 0.0001, *** p < 0.001, ** p<0.01, * p<0.05. Non-parametric One-way ANOVA (Kruskal Wallis) with Dunn’s posthoc test was used to analyze organ burden in mice.Two-way Anova was used to analyze the grouped data; p values **** p < 0.0001, *** p < 0.001, ** p<0.01, * p<0.05

STM WT invasion was reduced upon BF-8 treatment. In contrast, STM *ΔluxS* invasion increased with the treatment of DPD and STM WT spent media **(S4A)**. The immunofluorescence study with the histopathological sections of the mouse ileum further validated the reduced invasion and colonization of the mutants in the intestine **(Fig. 3D; S4B)**.

It is shown that the master regulator HilD regulates *Salmonella* invasion and motility via activating gene expression of the type 3 secretion system 1 (SPI-1) and motility genes(51, 52). We hypothesized that LuxS/AI-2 signaling regulates the *hilD* and other SPI-1 master regulators to modulate the invasion or infection in host cells. We observed that *hilD, hilA,* and *invF* master regulators of SPI-1 were downregulated in STM *ΔluxS*, STM *ΔlsrB*, and STM *ΔlsrK in vitro* LB media compared to STM WT **(Fig. 4 E-G)**. Thus, *Salmonella* requires LuxS/AI-2 signaling to invade successfully within the IECs *in vitro* and *in vivo* via regulating SPI-1 regulators.

**Fig. 4.**
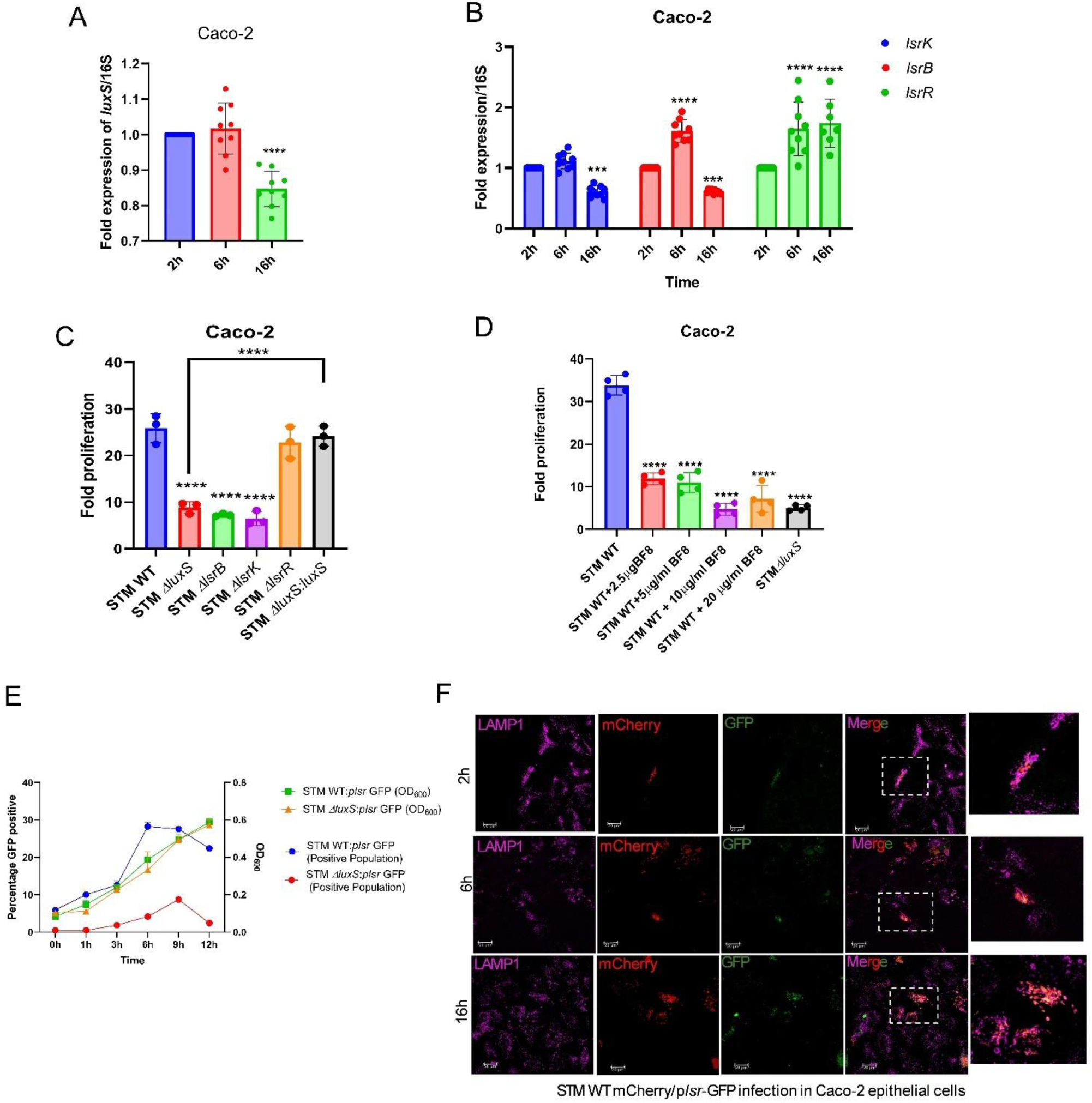
LuxS/AI-2 signaling is active and facilitates survival within Caco-2 epithelial cells. mRNA expression of **(A)** *luxS,* and **(B)** *lsr* operon genes *lsrB*, *lsrK,* and *lsrR* in STM WT upon infection in Caco-2 epithelial cells. Represented as Mean+/-SDof N=3, n=3. Fold proliferation of **(C)** STM WT and knockout strains, **(D)** STM WT treated with BF-8. Represented as Mean+/-SEM of N=3, n=3. **(E)** Quorum sensing reporter strains growth kinetics and flow cytometry analysis of GFP-positive bacterial cells. **(F)** Immunofluorescence microscopy analysis of STM WT quorum sensing reporter strains upon infection in Caco-2 epithelial cells. LAMP1 marker-*Salmonella* containing vacuole (SCV), mCherry-Constitutive expression, GFP-quorum sensing active STM. One-way ANOVA with Dunnet’s post-hoc test was used to analyze the data; p values **** p < 0.0001, *** p < 0.001, ** p<0.01, * p<0.05. Two-way Anova was used to analyze the grouped data; p values **** p < 0.0001, *** p < 0.001, ** p<0.01, * p<0.05.

### LuxS/AI-2 signaling is active and facilitates survival within Caco-2 epithelial cells

As the LuxS/AI-2 signaling is a key language for *Salmonella* communication,, we sought to explore its role in the next stages of infection. To begin, we asked how *Salmonella* Typhimurium regulates the expression of the genes within the *lsr* operon and *luxS* upon infection into host cells? We evaluated the mRNA expression of *luxS* and *lsr* operon genes (*lsrB*, *lsrK*, *lsrR*) in STM WT upon infection in the Caco2 cell line. We noted that the *lsr* operon genes were slightly higher at post 6h infection compared to 2h post-infection, and further, it downregulated at late 16h post-infection **(Fig. 4A)** while *luxS* expression was similar at 2h and 6h post-infection and downregulated at 16h post-infection compared to 2h **(Fig. 4B)**. Consistent expression of genes suggests that AI-2 signaling may be active inside the host cell. Following this, we observed that STM *ΔluxS*, STM *ΔlsrB*, and STM *ΔlsrK* proliferate less in comparison to STM WT, STM *ΔlsrR,* and *luxS* complemented STM *ΔluxS* within the Caco-2 cells **(Fig. 4C)**. The fold replication was reduced in the BF-8-treated culture of STM WT **(Fig. 4D)**. Conversely, upon treatment of WT spent media and exogenous DPD molecule, STM *ΔluxS* fold replication was recovered **(S5A, B)**. To further confirm whether this signaling is active inside the host cells, we generated the reporter strain, which produces green fluorescence on LuxS/AI-2 activation. Firstly, we checked the quorum sensing active bacteria during the *in vitro* growth in LB media. Flow cytometry and fluorescence microscopy results suggest that quorum-sensing active bacterial cells were increased in mid-log phase to early stationary phase **(Fig. 4E; S5C)**. Moreover, when we gave the STM WT spent media treatment to STM *ΔluxS* (Containing reporter construct), we observed that STM *ΔluxS* became quorum sensing active (**S5D)**. Upon infection in Caco-2 epithelial cells with reporter STM strain suggests that Lux/AI-2 signaling is active inside the host cells **(Fig. 4F)**. Taken together, this suggests that LuxS/AI-2 signaling is active and facilitates survival within Caco-2 epithelial cells.

### LuxS/AI-2 cascade facilitates *Salmonella* survival by evading host antimicrobial peptide defence mechanisms

The innate immune system within the gut relies substantially on antimicrobial peptides (AMPs) from epithelial cells, including cathelicidins and defensins. They help maintain intestinal homeostasis by targeting pathogens and regulating the gut microbiota(53). Our previous study showed that the LsrR repressor protein negatively modulates the expression of the *phoP* gene, which protects against acidic stress(54). Several studies revealed cationic antimicrobial peptides can activate the PhoP/PhoQ two-component system (TCS), and the PhoP/PhoQ two-component system TCS regulates the expression and activation of *pmrABD* and provides protection against the AMPs (55–62). We considered that LuxS/AI-2 signaling might aid in combating antimicrobial peptide sensitivity via regulating *phoP/phoQ*. In continuation, we observed that STM *ΔluxS*, STM *ΔlsrB*, and STM *ΔlsrK* were more sensitive compared to STM WT and STM *ΔlsrR* under *in vitro* conditions in the presence of sub-MIC concentration of polymyxin **(Fig. 5A; S 6A, B)**. Subsequently, we found that the mRNA expression levels of *pmrD*, *pmrA*, and *pmrB* were reduced in STM *ΔluxS*, STM *ΔlsrB*, and STM *ΔlsrK* compared to STM WT under polymyxin-treated conditions *in vitro* in LB media **(Fig. 5B)**. However, in STM *ΔlsrR*, their expression levels were comparable to those in STM WT. Thereafter, *pmrD*, *pmrA*, and *pmrB* gene expression level reduction was observed upon infection in Caco-2 epithelial cells in STM *ΔluxS*, STM *ΔlsrB*, and STM *ΔlsrK* compared to STM WT **(Fig. 5 C-E)**. Conclusively, LuxS/AI-2 signaling facilitates survival within Caco-2 epithelial cells by regulating the *pmrD/AB* gene expression.

**Fig. 5.**
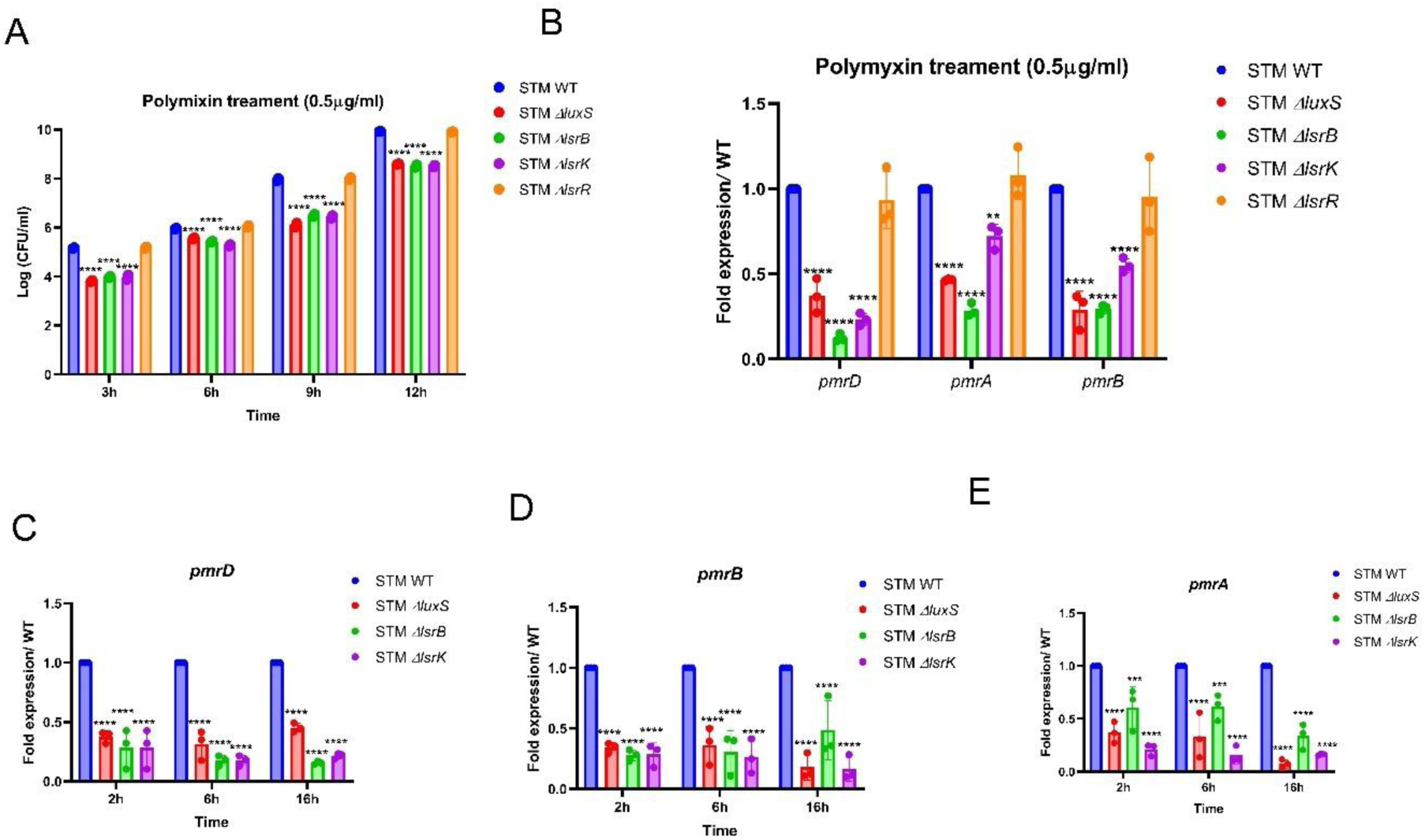
LuxS/AI-2 cascade facilitates *Salmonella* survival by evading host antimicrobial peptide defence mechanisms. **(A)** *In vitro* polymyxin treatment survival assay of STM WT, STM *ΔluxS*, STM *ΔlsrB*, STM *ΔlsrK,* and STM *ΔlsrR.* Represented as Mean+/-SD of N=3, n=2. **(B)** mRNA expression of *pmrD*, *pmrA,* and *pmrB* in STM WT, STM *ΔluxS*, STM *ΔlsrB*, STM *ΔlsrK,* and STM *ΔlsrR* upon treatment of polymyxin in vitro LB media in mid-log phase culture (6h). Represented as Mean+/-SD of N=3, n=3. mRNA expression of **(C)** *pmrD*, **(D)** *pmrA* and **(E)** *pmrB* in STM WT, STM *ΔluxS*, STM *ΔlsrB*, STM *ΔlsrK,* and STM *ΔlsrR* upon infection in Caco-2 epithelial cells. Represented as Mean+/-SD of N=3, n=3. Two-way Anova was used to analyze the grouped data; p values **** p < 0.0001, *** p < 0.001, ** p<0.01, * p<0.05

### LuxS/AI-2 signaling regulates critical gene targets to assist in multiple stages of *Salmonella* infection in the host

As we observed, LuxS/AI-2 signaling is involved in various stages of *Salmonella* infection by regulating multiple genes. Following this, we performed transcriptomic analysis between STM WT and quorum sensing knockout strains STM *ΔluxS*, STM *ΔlsrB, and* STM *ΔlsrK.* We figured out that over the 4728 gene variables, 507, 1291, and 225 genes were significantly differentially expressed in STM *ΔluxS*, STM *ΔlsrB,* and STM *ΔlsrK,* respectively. Out of which, 316 are down-regulated and 191 are up-regulated in STM *ΔluxS,* 902 are down-regulated and 389 genes are up-regulated in STM *ΔlsrB,* while 106 are down-regulated and 119 are up-regulated in STM *ΔlsrK* in comparison to STM WT **(Fig. 6A-C)**. Gene enrichment and KEGG analysis revealed that down-regulated genes belong to the flagellar classes (*fliC*/*fljB* family gene), chemotaxis (*cheY*, *motA*, *motB* etc.), secretory protein and virulence-associated genes (*sopD2*, *sseI*, *pipB*, *sopB*, *sifA*) etc **(Fig. 6D, E)**. Interestingly, the heatmap of the top 50 genes also includes genes related to flagellar motility *fliC*/*fljB*, chemotaxis gene *motA,* and virulence genes (*hilD*, *pipB*, *sseI,* etc.) **(S7A-C)**. Taken together, the transcriptomic analysis suggests that the Lux/AI-2 signaling regulates multiple critical gene targets to assist in multiple stages of *Salmonella* infection in the host.

**Fig. 6.**
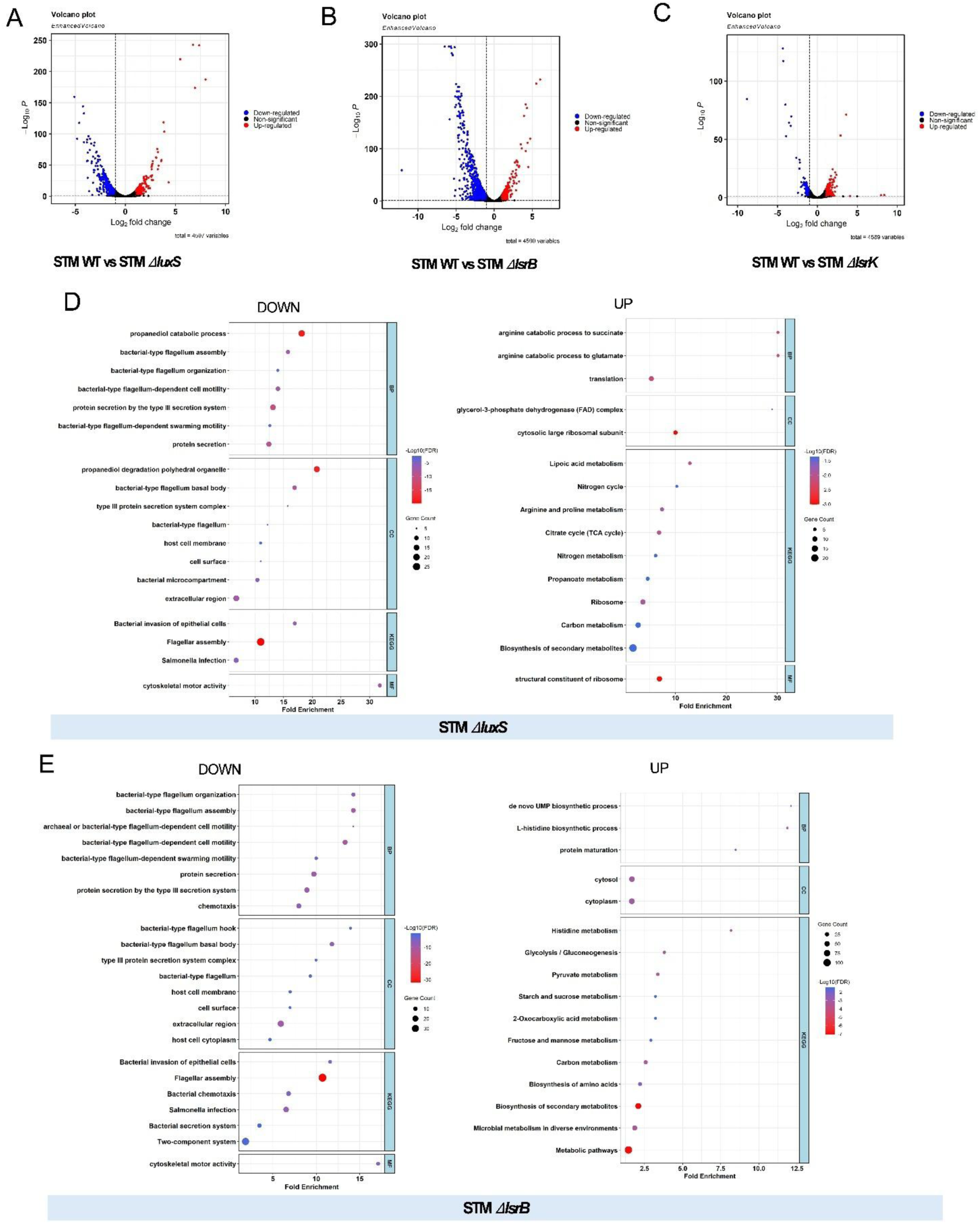
LuxS/AI-2 signaling regulates critical gene targets to assist in multiple stages of *Salmonella* infection in the host. Transcriptomic analysis of STM WT, STM *ΔluxS*, STM *ΔlsrB*, and STM *ΔlsrK* grown in LB media **(A-C)**. Gene enrichment (GO) and KEGG analysis in STM *ΔluxS* and STM *ΔlsrB* **(D-E).**

### Inhibition of LuxS/AI-2 signaling with antibiotics reduces *Salmonella* colonization *in vivo* and enhances mice survival

*Salmonella* can mediate its uptake into enterocytes by delivering effector molecules through Type 3 secretion systems (T3SS). It can also be phagocytosed by immune cells such as macrophages, dendritic cells, and neutrophils residing in Peyer’s patches. However, *Salmonella* preferentially enters specialized cells called microfold cells or M-cells of the Peyer’s patches in the intestinal epithelium. After breaching the epithelial cells at the Peyer’s patches in the distal ileum, it disseminates to the liver, spleen, and mesenteric lymph node (MLN), the secondary infection sites(63, 64). *Salmonella* is known to be a successful pathogen that can overcome various host immune responses such as oxidative stress, nitrosative stress, acidic pH, etc(65). In our recent study, STM *ΔluxS*, STM *ΔlsrB*, and STM *ΔlsrK* showed less colonization and delayed mouse mortality compared to STM WT infection, and upon treatment of BF-8 (4mg/kg mice body weight) to STM WT-infected mice, organ burden was reduced, and mouse survivability increased(54). Following this, we observed in competition between STM WT with STM *ΔluxS* or STM *ΔlsrB*, or STM *ΔlsrK* in mice co-infection studies, STM WT outcompeted all three mutants in the liver, spleen, MLN, and intestine **(Fig. 7A, B)**. Further, we questioned whether, upon treatment with BF-8 in combination with low-concentration doses (2mg/kg of each) of antibiotics ciprofloxacin and ceftazidime, the infection and mortality of mice could be cured. So, we first infected the mice with STM WT by oral gavage and started the treatment on the 3rd day post-infection (Fig. 7C). On the 3rd day post-infection, we observed organ colonization and slight weight reduction, confirming infection **(S8 A-C)**. After each day of treatment of BF-8 with or without antibiotics for consecutive days, on the 6^th^ day post-infection, we noted that organ colonization was significantly less with the combination of low concentration 2mg/kg of body weight (of BF-8 and ciprofloxacin or ceftazidime **(Fig. 7C-F; S8D, E)**. With the treatment of three doses of a combination of BF-8 with ciprofloxacin or ceftazidime, started on the 3^rd^ day post-infection, mice survived till 14 or 15 days, respectively. On the other hand, low doses of BF-8 and antibiotics also delayed the mortality of mice compared to the untreated mice **(Fig. 7G, H; S8 F)**. Furthermore, tissue histopathology of liver and spleen, suggested that a combination treatment of BF-8 inhibitor with antibiotics improved the health of the mice more effectively with a less disease score (**Fig. 7I; S9)**. These data suggest that the AI-2 mediated signaling is required for *in vivo* colonization of *Salmonella* in mice, and signaling inhibition can be a promising strategy in combination with low doses of antibiotics for treatment of *Salmonella* infection in the era of emerging antibiotic-resistant *Salmonella* strains.

**Fig. 7.**
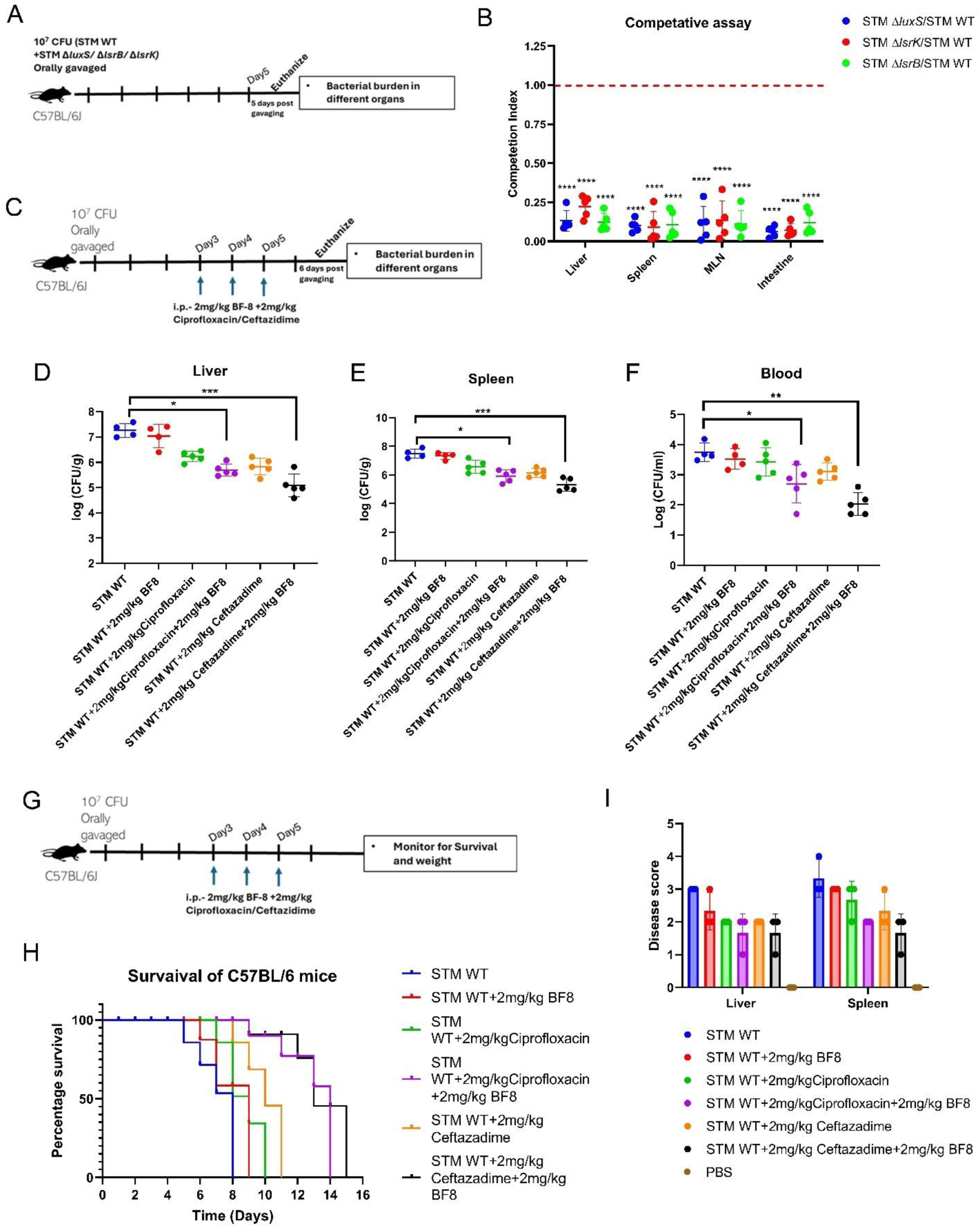
Inhibition of LuxS/AI-2 signaling with antibiotics reduces *Salmonella* colonization *in vivo* and enhances mice survival. **(A)** The Competitive experimental protocol for organ burden in C57BL/6J mice by orally gavaging a total of 10^7^ CFU per mouse. **(B)**The organ burden post 5 days of oral gavage in a competition assay. Represented as Mean+/-SD of N=2, n=5 **(C)**. The experimental protocol for C57BL/6J mice infected with STM WT by orally gavaging 10^7^ and BF-8 and an antibiotic treatment. Organ burden post treatment **(D)** Liver, (**E)** Spleen, and **(F)** Blood. Represented as Mean+/-SD of N=2, n=5. **(G)** The experimental protocol of mouse survival in C57BL/6J mice involved orally gavaging 10^7^ CFU per mouse upon treatment with the BF-8 inhibitor and a combination of antibiotics (through the Intraperitoneal route). **(H)** Percentage survival of Mice upon infection with STM WT with and without treatment. **(I)** Disease score from the histopathological sections of the liver and spleen. Non-parametric One-way ANOVA (Kruskal-Wallis) with Dunn’s post hoc test was used to analyze organ burden in mice.

## Discussion

Adapting its tactics and employing multiple defense systems, *Salmonella* has become a potent enteric pathogen, occupying several host niches during its pathogenesis. The opening moves during its pathogenesis involve the expansion and competition in the gut, followed by attachment and invasion. To do so, *Salmonella* must first reach the site of infection. Of the several factors, chemotaxis and motility contribute majorly to such a process. Recently, it was shown that AI-2, a quorum-sensing molecule, can work as a chemoattractant and help in aggregation(24). Here, we showed that *Salmonella* LuxS/AI-2 signaling controls motility via regulating motility, flagellar gene expression, and flagellar organisation. *Salmonella* produces and releases the AI-2 molecules, which, upon reaching an optimal concentration in the environment, begin to accumulate intracellularly via the ABC transporter. The phosphorylation of the intracellular AI-2 by LsrK is crucial for its function and orchestration of its gene targets (66). Since the accumulation of AI-2 upregulates the *lsr* operon, we also observe that the genes *lsrB*, *lsrK*, and *lsrR* were upregulated during the mid-log and late-log phases(54). This provides an additional context to our current observations, as flagella formation is primarily during its exponential growth phases *in vitro*.

The gastrointestinal tract is a dynamic environment where microbes compete for space and nutrients through various mechanisms. Many gut-associated bacteria produce AI-2 via the LuxS enzyme, including a significant proportion of Firmicutes and Proteobacteria, and a few members of the phyla Bacteroidetes and Actinobacteria. (67, 68). Besides, species of the families Barnesiellaceae and Muribaculaceae also contribute to AI-2 production in the gut (69). Being an enteric pathogen and a sensor of the universal communication signal, *Salmonella* may profit from the AI-2 produced by the gut commensal in the intestinal lumen. Through its ABC transporter, *Salmonella* may readily take up the extracellular AI-2 produced by the gut microbes. This further demonstrates that intra-bacterial communication and interactions might be exploited by pathogens to win the arms race.

Natural fluid flow in the intestine can remove autoinducers, disrupting quorum sensing. *Staphylococcus aureus* utilizes autoinducers in the crypts (non-existent flow) and enhances pathogenicity by producing enterotoxin B (45). Similarly, *Salmonella* targets the ileum crypts to invade epithelial cells, ensuring survival and replication. Natural dynamics are crucial in bacterial adaptation and virulence(48). Motility towards the ileum of the intestine is essential, and sensing AI-2 plays a pivotal role in *Salmonella* pathogenesis. Our results suggest that *Salmonella* can move toward the AI-2 molecule, and LsrB might work as a chemoreceptor. As previously studied, LsrB in *E.coli* acts as a receptor for AI-2 and aids in the sensing and drives chemotaxis towards the AI-2 pool (70). Similarly, we speculate that in *Salmonella,* the LsrB receptor also assists in sensing AI-2, while the AI-2 is internalised through the ABC receptor, which tunes the expression of chemotaxis and motility gene expression to drive motion towards the AI-2 pool. HilD is the master regulator of motility and the SPI-1 gene. The present study shows that LuxS/AI-2 signaling regulates the *hilD* gene expression, thereby aiding in motility, attachment, and invasion of epithelial cells *in vitro* and *in vivo* conditions. Additionally, we observe that the SPI-1 genes *hilA* and *invF* are also underexpressed in the quorum-sensing mutants. This likely accounts for the impaired adhesion and invasion of these mutants, as the SPI-1 encoded T3SS system and effector molecule mediate host cell invasion.

Our findings further indicated that accumulation and phosphorylation of AI-2 are necessary for controlling the expression of the *pmrD/AB*. The *pmrD/AB* could be regulated by PhoP, resulting in reduced survival of STM *ΔluxS*, STM *ΔlsrB*, and STM *ΔlsrK in vitro* and in the epithelial cells (71, 72). Our previous study highlighted the regulatory mechanism of LuxS/AI-2 in controlling the *phoP/Q* transcription (54). Hence, in such a context, our present study further highlights the broader, interconnected regulatory roles of LuxS/AI-2 in several defence strategies of this pathogen. Transcriptomic analysis revealed that LuxS/AI-2, the language of *Salmonella,* can control various critical genes in *Salmonella.* These genes are involved in flagellar motility, invasion, cell surface and membrane functions, and secretory pathways. This underscores how the quorum-sensing cascade regulates numerous genetic programs that are involved in the various stages of *Salmonella* infection. As previously reported, the regulation of the *phoP/Q* operon by the LsrR repressor through the promoter binding activity of the repressor, it can be assumed that some of these genetic controls may also involve the LsrR binding activity (54). Our present work also demonstrated that the organ burden of *Salmonella* was reduced upon treatment of mice with inhibitor BF-8 in combination with low doses of antibiotics ciprofloxacin and ceftazidime. Considering the increasing emergence of antimicrobial-resistant *Salmonella* strains worldwide, there is a need for alternative approaches to curb infection. As we point out to the broader and interconnected regulatory mechanisms by the LuxS/AI-2 signaling in *Salmonella*, intervening in such a critical signaling process proves to be a promising infection resolution strategy. Considering the pervasive presence of AI-2, it appears paradoxical that *Salmonella* cells depend on it to colonize a specific niche within the gut. One plausible rationale for this might be that AI-2 chemotaxis becomes significant under dysbiosis conditions that suppress microbiota and promote the proliferation of *Enterobacteriaceae* (73, 74). Our study provides evidence for a causal link between AI-2 chemotaxis, gut colonization, and niche segregation of *Salmonella* strains in the murine gut. It further establishes a link between AI-2 chemotaxis, invasion, and colonization *in vivo*. We suggest that similar mechanisms of AI-2-mediated host colonization might exist in other chemotactic enteric bacteria, and inhibition of LuxS/AI-2 signaling might open avenues for antibiotic resistance of pathogenic bacteria.

## Data availability

The RNA-seq data generated in this study have been deposited in NCBI’s Gene Expression Omnibus (GEO) and can be accessed with the GEO series accession numbers GSE305594. All the data supporting the conclusions of this study are available in the paper and supplemental materials.

## Ethics statement

All experiments comply with the rules set forth by the Indian Institute of Science, Bangalore’s IEAC. The approved protocol number is CAF/ Ethics/852/2021. The Committee for Control and Supervision of Experiments on Animals (CPCSEA), a statutory committee established under Chapter 4, Section 15 (1) of the Prevention of Cruelty to Animals Act 1960, and National Animal Care provided guidelines that were meticulously adhered to during all animal experiments, all of which were approved by the Institutional Animal Ethics Committee. (Registration No. 435 48/1999/ CPCSEA). The Institutional Animal Ethics Committee approved every animal experiment, and the National Animal Care Guidelines were meticulously followed.

## Author contribution

AS and DC conceived the study. AS and DC designed experiments. AS performed experiments. AS and AVN performed an electron microscopy experiment. RSR analyzed the tissue histopathology. AS analyzed the data, prepared the figures, and drafted the manuscript. AS, AVN, and DC reviewed and edited the manuscript. DC supervised the work. All the authors read and approved the manuscript.

## Acknowledgment

The Departmental and Divisional Confocal Facility, Departmental Real-Time PCR Facility, Divisional Electron Microscopy facility, and Central Animal Facility at IISc are duly acknowledged. Mr. Arjun and Ms. Syama are acknowledged for their help with image acquisition. Ms. Kirtan is acknowledged for her help with Electron microscope image acquisition. Ms. Aneesha and Ms. Nanditha are acknowledged for their technical help.

