## Supplementary Information for "Interbacterial AI-2 communication drives stage-specific genetic programs to support *Salmonella* colonisation in the murine gut"

**S1:**

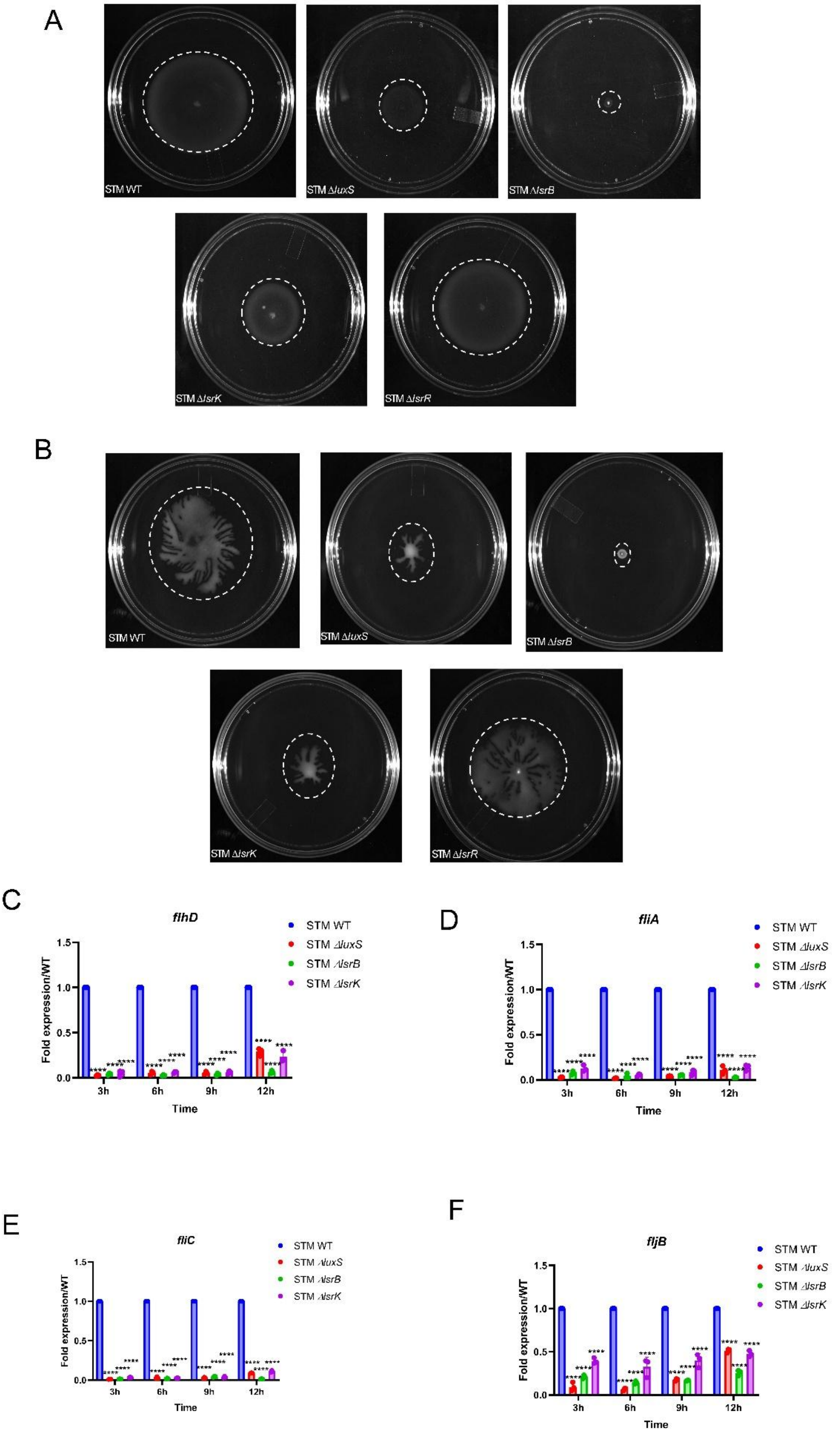
A
STM WT
STM ΔluxS
STM ΔlsrB
STM ΔlsrK
STM ΔlsrR
B
STM WT
STM ΔluxS
STM ΔlsrB
STM ΔlsrK
STM ΔlsrR
C
flhD
Fold expression/WT
Time
3h
6h
9h
12h
STM WT
STM ΔluxS
STM ΔlsrB
STM ΔlsrK
D
fliA
Fold expression/WT
Time
STM WT
STM ΔluxS
STM ΔlsrB
STM ΔlsrK
E
fliC
Fold expression/WT
Time
STM WT
STM ΔluxS
STM ΔlsrB
STM ΔlsrK
F
fljB
Fold expression/WT
Time
STM WT
STM ΔluxS
STM ΔlsrB
STM ΔlsrK

**S1: (A)** Swim motility assay, **(B)** Swarm agar assay; of STM WT, STM *ΔluxS*, STM *ΔlsrB*, STM *ΔlsrK,* and STM *ΔlsrR.* Representative image of N=3, n=3,**:** mRNA expression of **(C)** *flhD,* **(D)** *fliA,* **(E)** *fliC,* and **(F)** *fljB,* in STM WT, STM *ΔluxS*, STM *ΔlsrB*, STM *ΔlsrK,* and STM *ΔlsrR* in *in vitro* LB media. Represented as Mean+/-SD of N=3, n=3, Two-wa Anova was used to analyze the grouped data; p values **** p < 0.0001, *** p < 0.001, ** p<0.01, * p<0.05

**S2:**

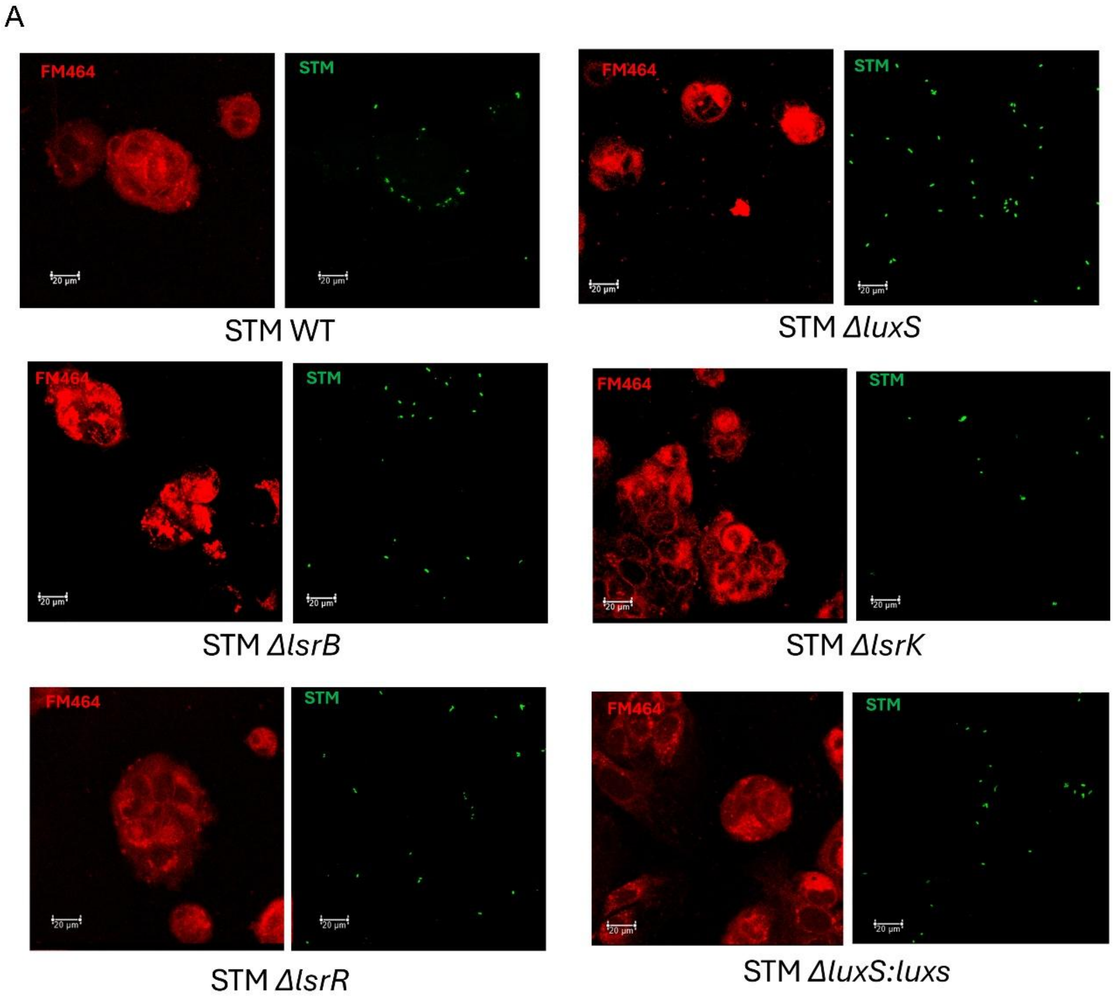

**S2: (A)** Fluorescence microscopy of adhesion assay in Caco-2 epithelial cells. Red- FM464 dye for Caco-2 cells, Green- STM strains. Representative image of N=3, n>10.

## S3:

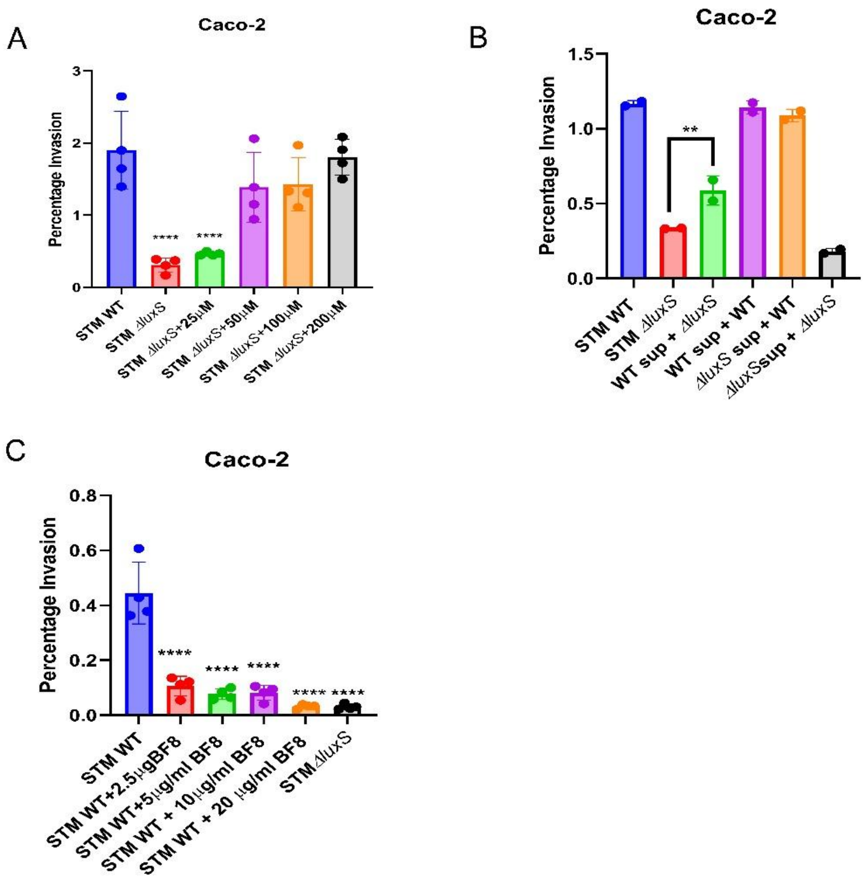


**S3:** Percentage invasion in Caco-2 epithelial cells **(A)** upon treatment of exogenous DPD molecule to STM *ΔluxS*, **(B)** Upon treatment of spent media (STM WT) to STM *ΔluxS,* **(C)** Upon treatment of BF-8 to STM WT**.** Represented as Mean+/-SD of $N=3$, $n=2$.One-way ANOVA with Dunnet's post-hoc test was used to analyze the data; p values **** $p < 0.0001$, *** $p < 0.001$, ** $p<0.01$, * $p<0.05$.

## S4:

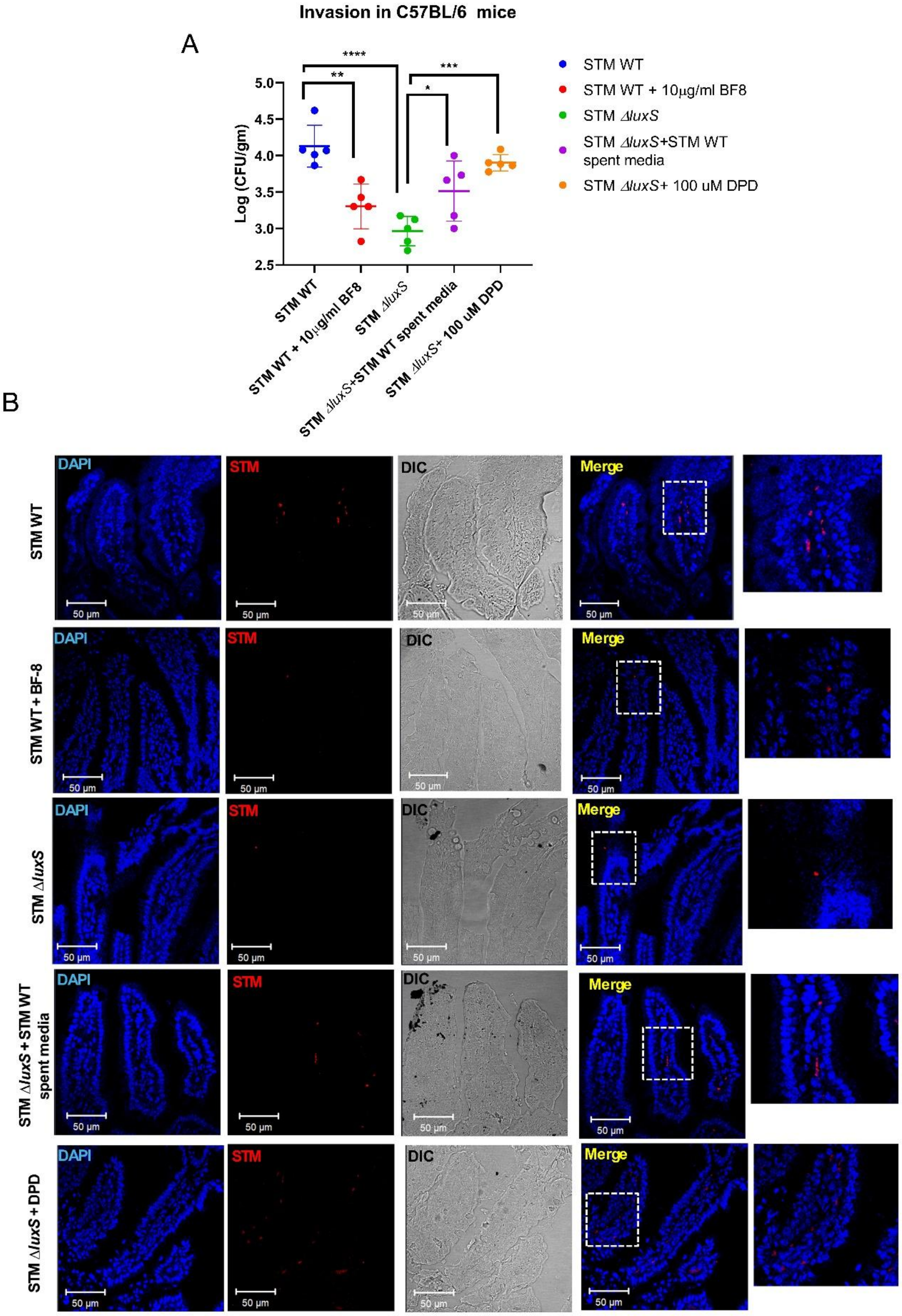

Invasion in C57BL/6 mice
A
Log (CFU/gm)
5.0
4.5
4.0
3.5
3.0
2.5
****
**
***
*
STM WT
STM WT + 10µg/ml BF8
STM ΔluxS
STM ΔluxS+STM WT spent media
STM ΔluxS+ 100 uM DPD
B
DAPI
STM
DIC
Merge
50 µm
STM WT
STM WT + BF-8
STM ΔluxS
STM ΔluxS + STM WT spent media
STM ΔluxS + DPD

**S4: (A)** STM WT treated with BF-8, STM *ΔluxS* treated with STM WT spent media and DPD molecule, followed by invasion assay in mice; organ burden in the intestine post 6h of infection. Represented as Mean+/-SD of N=2, n=5.**(B)** Immunofluorescence of histopathological sections of mice intestine (Peyer's patches) to study invasion, DAPI is used to stain the nucleic acids in the cells, and Anti-Salmonella (LPS) (Cy3 tagged secondary antibody used-Red). Non-parametric One-way ANOVA (Kruskal Wallis) with Dunn's posthoc test was used to analyze organ burden in mice.

# S5:

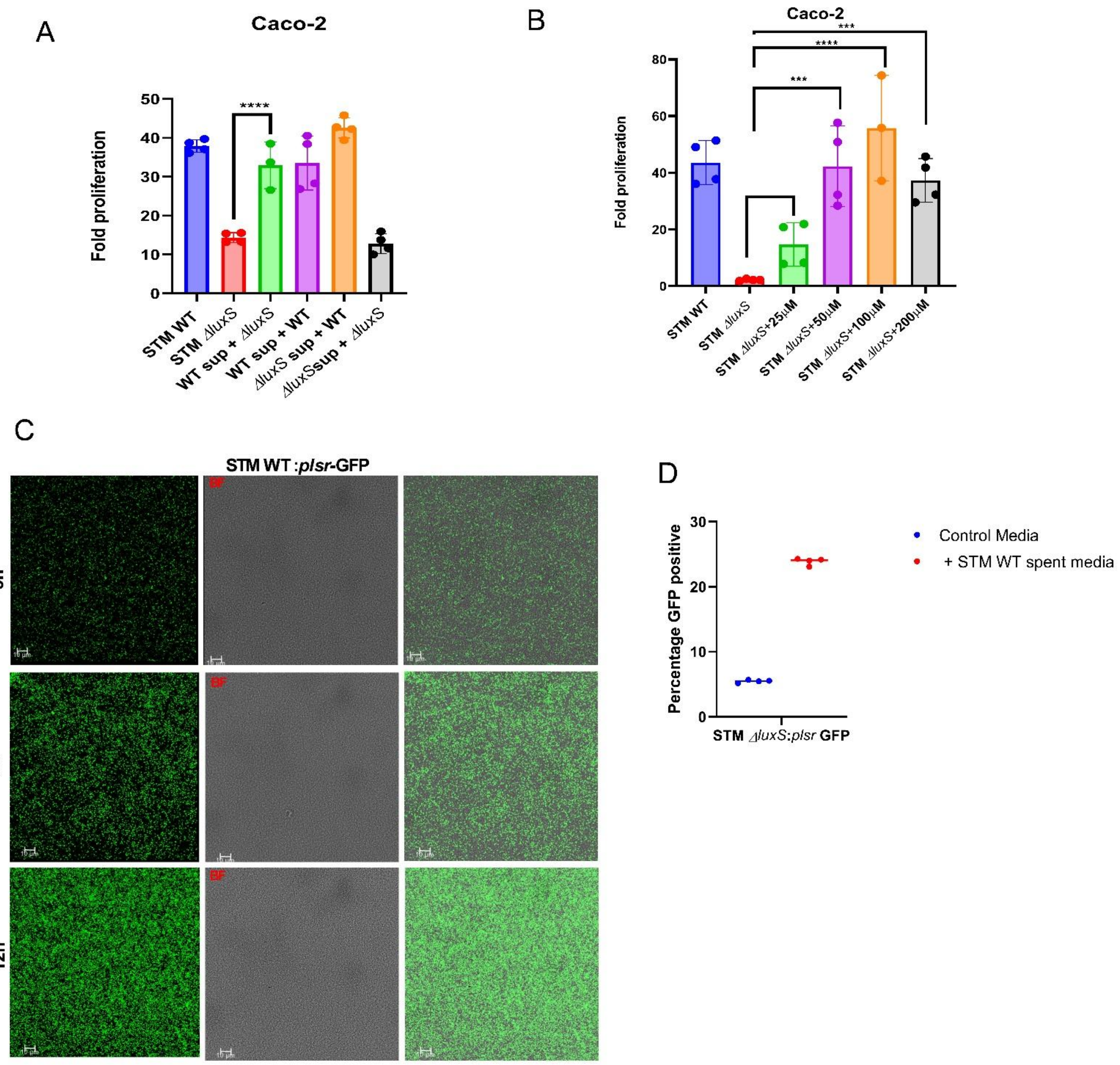


**S5:** Fold proliferation of **(A)** STM *ΔluxS* upon treatment with STM WT spent media, **(B)** STM *ΔluxS* upon treatment with DPD molecule. Represented as Mean+/-SD of N=3, n=2. **(C)** STM WT: *plsr*-GFP reporter strains grown in LB media, and immunofluorescence microscopy was done for confirmation of LuxS/AI-2 signaling active cells. **(D)** Flow cytometry analysis of STM *ΔluxS* :*plsr*-GFP upon treatment of STM WT spent media. One-way ANOVA with Dunnet's post-hoc test was used to analyze the data; p values **** $p < 0.0001$, *** $p < 0.001$, ** $p<0.01$, * $p<0.05$.

## S6:

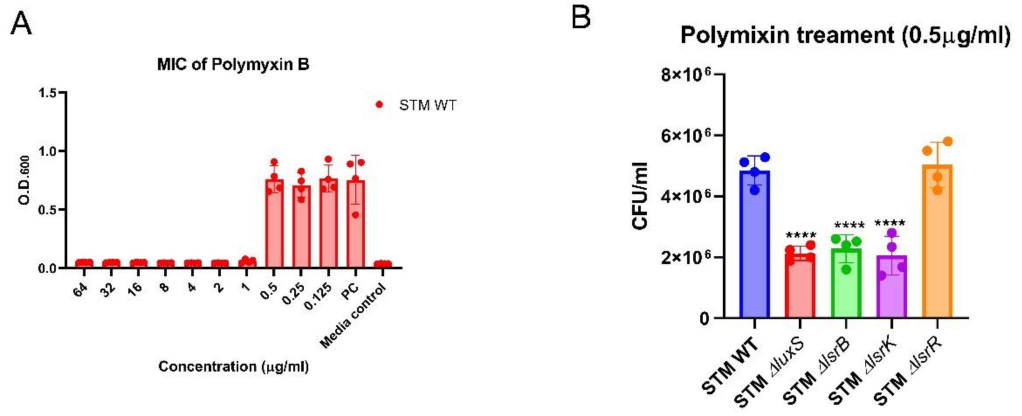


**S6: (A)** Minimum inhibitory concentration of polymyxin for STM WT. Represented as Mean+/-SD of N=2, n=4. **(B)** Survival assay of STM WT, STM *ΔluxS*, STM *ΔlsrB*, STM *ΔlsrK,* and STM *ΔlsrR in vitro* LB media upon treatment with sub-MIC concentration of polymyxin (2h incubation of treatment). Represented as Mean+/-SD of N=3, n=2. One-way ANOVA with Dunnet's post-hoc test was used to analyze the data; p values **** $p < 0.0001$, *** $p < 0.001$, ** $p<0.01$, * $p<0.05$.

## S7:

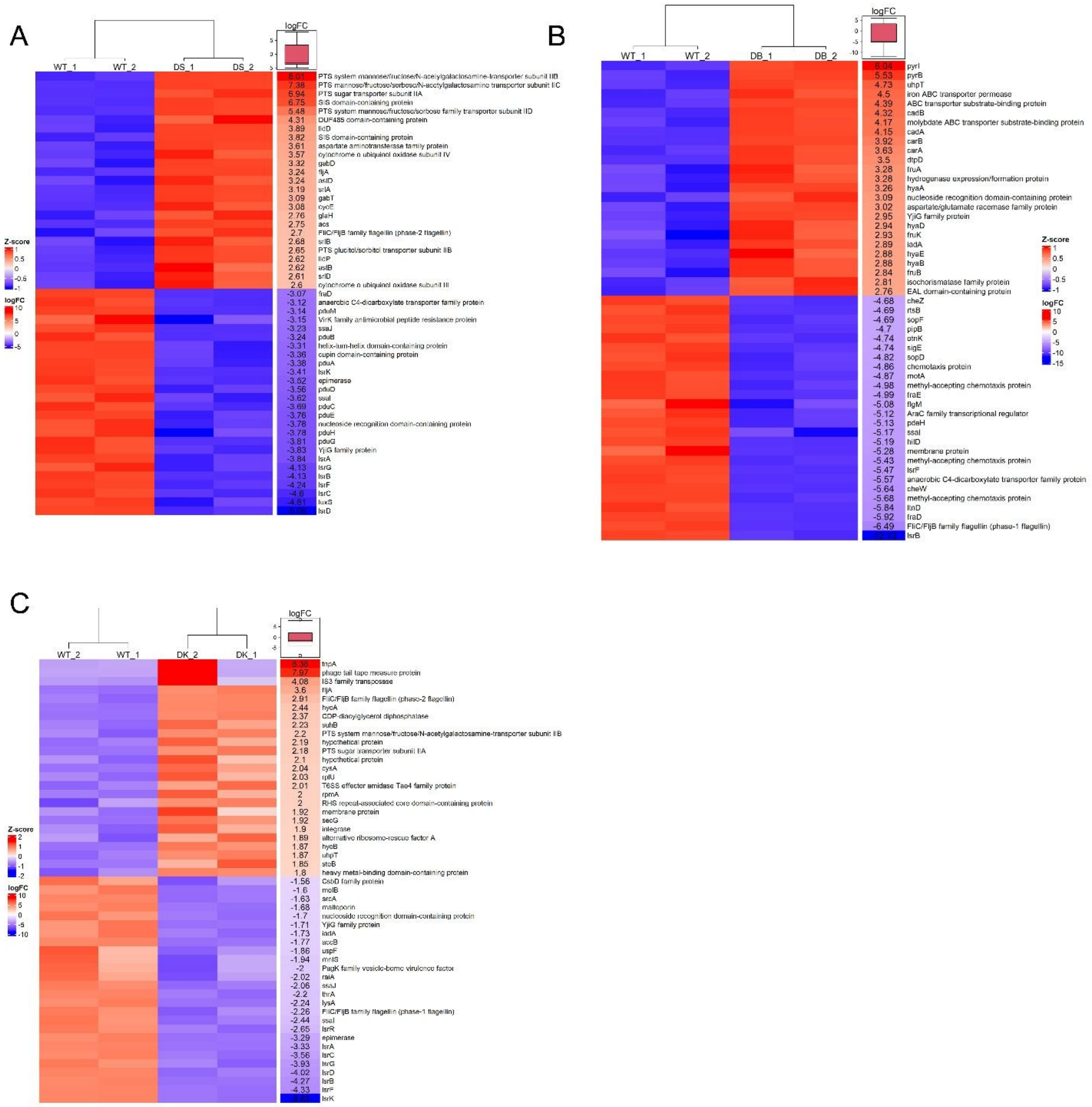


**S7:** Heat map of top 50 significantly differentially expressed genes in STM *ΔluxS*, STM *ΔlsrB*, and STM *ΔlsrK* in comparison with STM WT.

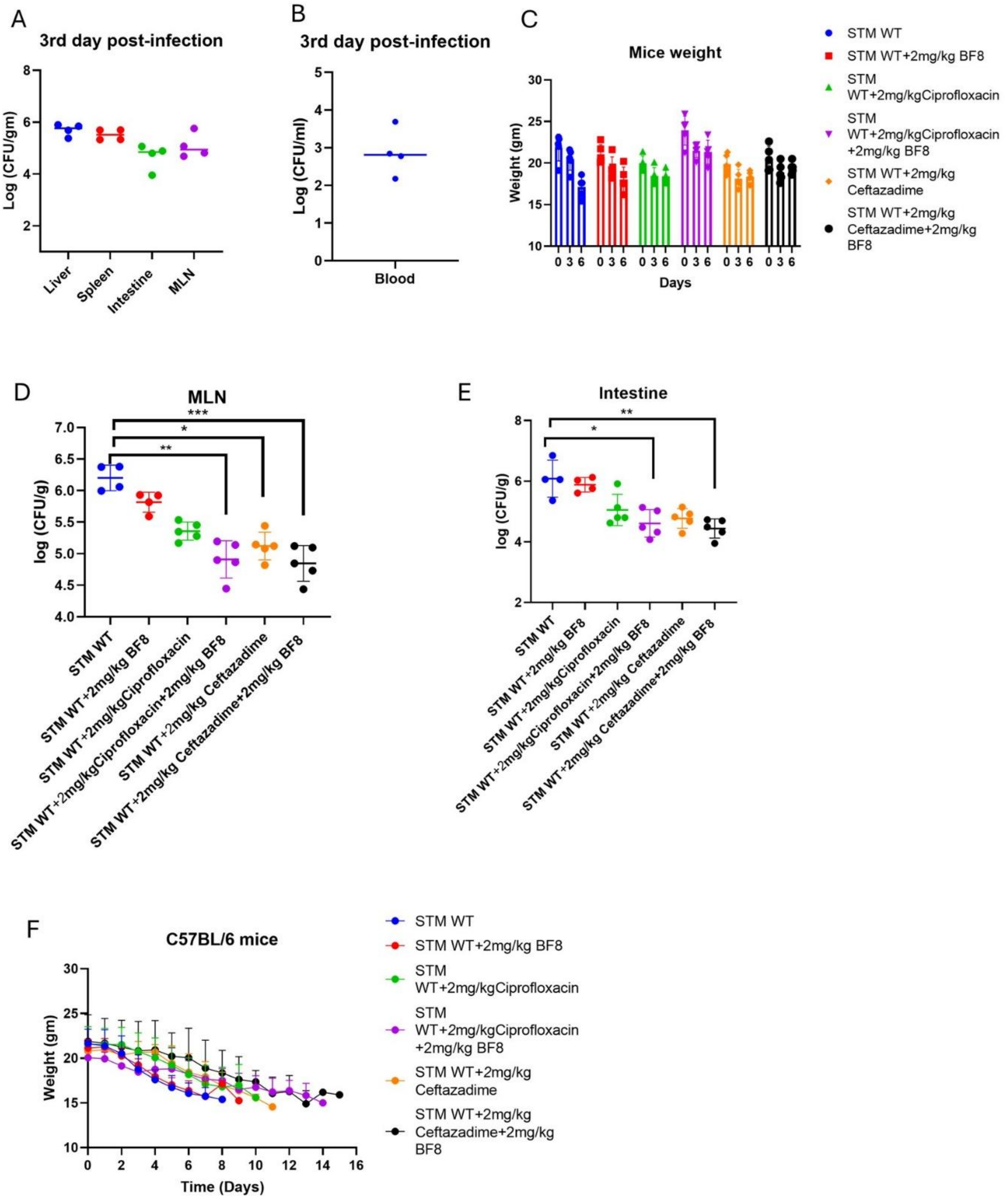
A
3rd day post-infection
Log (CFU/gm)
Liver
Spleen
Intestine
MLN
B
3rd day post-infection
Log (CFU/ml)
Blood
C
Mice weight
Weight (gm)
Days
STM WT
STM WT+2mg/kg BF8
STM WT+2mg/kgCiprofloxacin
STM WT+2mg/kgCiprofloxacin +2mg/kg BF8
STM WT+2mg/kg Ceftazadime
STM WT+2mg/kg Ceftazadime+2mg/kg BF8
D
MLN
log (CFU/g)
***
*
**
STM WT
STM WT+2mg/kg BF8
STM WT+2mg/kgCiprofloxacin
STM WT+2mg/kgCiprofloxacin+2mg/kg BF8
STM WT+2mg/kg Ceftazadime
STM WT+2mg/kg Ceftazadime+2mg/kg BF8
E
Intestine
log (CFU/g)
**
*
STM WT
STM WT+2mg/kg BF8
STM WT+2mg/kgCiprofloxacin
STM WT+2mg/kgCiprofloxacin+2mg/kg BF8
STM WT+2mg/kg Ceftazadime
STM WT+2mg/kg Ceftazadime+2mg/kg BF8
F
C57BL/6 mice
Weight (gm)
Time (Days)
STM WT
STM WT+2mg/kg BF8
STM WT+2mg/kgCiprofloxacin
STM WT+2mg/kgCiprofloxacin +2mg/kg BF8
STM WT+2mg/kg Ceftazadime
STM WT+2mg/kg Ceftazadime+2mg/kg BF8

**S8: (A, B)** Organ burden and Blood dissemination of STM WT infection post-3rd infection. **(C)** Weight measurement of mice post-3rd day of infection and post BF-8 and antibiotic treatment**.** C57BL/6J mice infected with STM WT by orally gavaging $10^7$ and BF-8 and an antibiotic treatment. Organ burden post treatment **(D)**MLN, (**E)** Intestine. Represented as Mean+/-SD of N=2, n=5. **(F)** Mice weight measurement, which were orally gavaged $10^7$ CFU per mouse, upon treatment with the BF-8 inhibitor and a combination of antibiotics (through the Intraperitoneal route). Non-parametric One-way ANOVA (Kruskal-Wallis) with Dunn's post hoc test was used to analyze organ burden in mice.

S9:

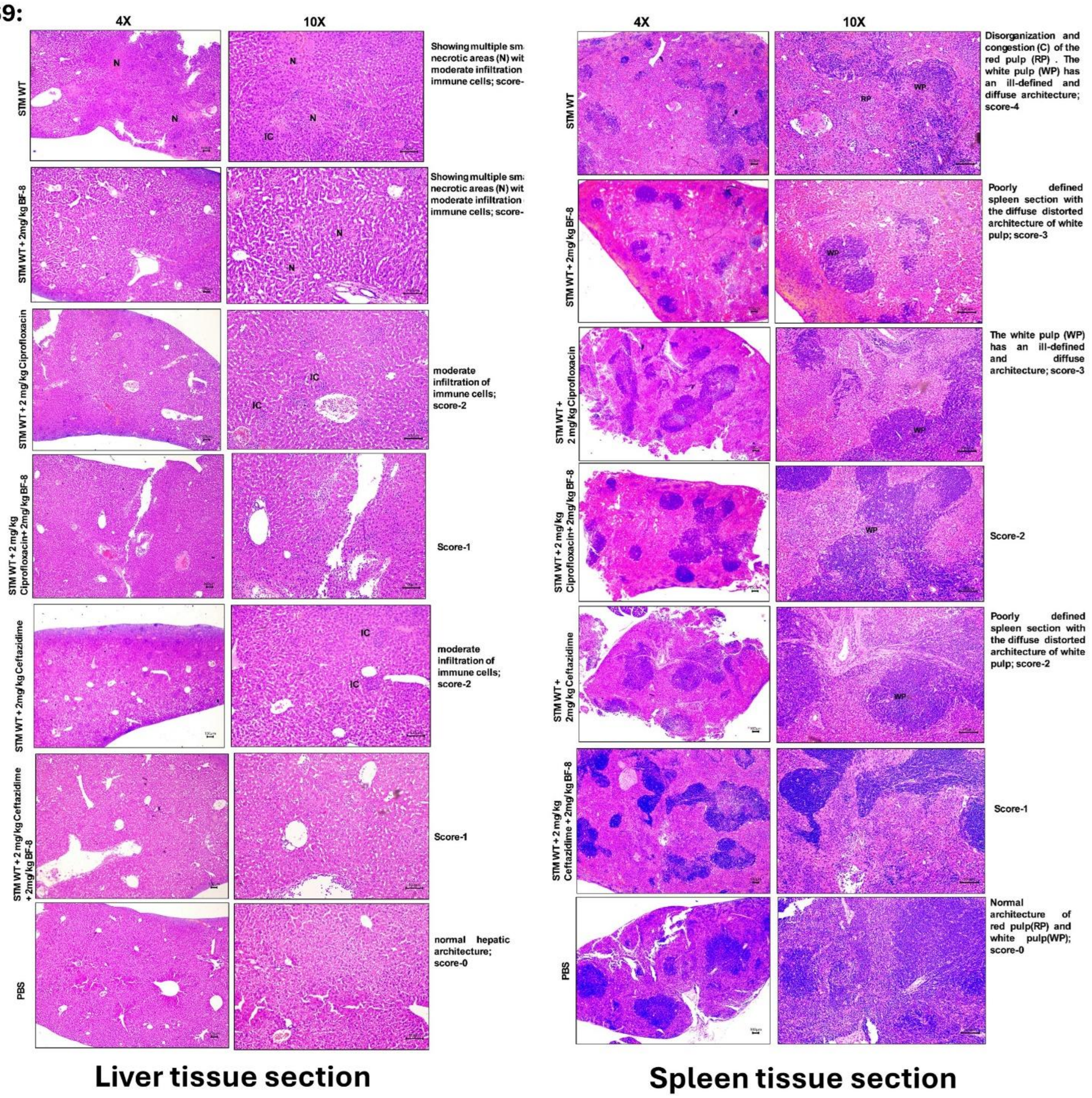


**S9: Histopathological scoring of hepatic necroinflammation and Spleen:** The hepatic necroinflammation score is done according to the criteria of the histological activity index (HAI) (Ishak et al. 1995) with little modification. The scoring of necroinflammation is graded as 0-4; for each of the pathological lesions (Severe -4 ; Moderate -3; Mild -2; Minor/minimum -1; No pathology -

0), considering the following pathological lesions: infiltration of immune cells, focal inflammation, and focal necrosis. Spleen section is done with disorganization and congestion (C) of the red pulp (RP) and the white pulp (WP) architecture. The scoring of the pathological changes is graded as 0-4; for each of the pathological lesions (Severe/ marked -4; Moderate -3; mild -2; Minor/minimum -1; No pathology -0).

**S Table 1** Bacterial strains and plasmid used in this study

| Strains/Plasmids | Characteristics | Sources |
|---|---|---|
| STM WT | No antibiotics | Gift from Prof. M. Hensel |
| STM $\Delta$*luxS* | Chl$^R$ | Previous study<br>https://doi.org/10.1101/2025.03.26.645541 |
| STM $\Delta$*lsrB* | Kan$^R$ | Previous study<br>https://doi.org/10.1101/2025.03.26.645541 |
| STM $\Delta$*lsrK* | Chl$^R$ | Previous study<br>https://doi.org/10.1101/2025.03.26.645541 |
| STM $\Delta$*lsrR* | Kan$^R$ | Previous study<br>https://doi.org/10.1101/2025.03.26.645541 |
| STM $\Delta$*luxS:luxS* | Chl$^R$, Amp$^R$<br>(complemented in pQE60 plasmid) | Previous study<br>https://doi.org/10.1101/2025.03.26.645541 |
| pKD3 plasmid | Chl$^R$ resistance cassette | Laboratory stock |
| pKD4 plasmid | Kan$^R$ resistance cassette | Laboratory stock |
| pKD46 plasmid | Plasmid expressing $\lambda$-red recombinase system, Amp$^R$ | Laboratory stock |

**S Table 2 Primers used in this study:**

| Primers | Sequence 5'…….3' |
|---|---|
| *luxS* knockout forward primer | CGGAGGTGACTAAATGCCATTATTAGATAGCTTCGCAGTC**CATATGAATATCCTCCTTAG** |
| *luxS* knockout reverse primer | CTGGAACCGCTTACAAATAAGACTAAATATGCAGTTCCTG**GTGTAGGCTGGAGCTGCTTC** |
| *luxS* knockout confirmation forward primer | GCAAAACACGCCTGACCCAA |
| *luxS* knockout confirmation reverse primer | CAATACACTCTGGCATCGTG |
| *luxS* expression forward primer | GCTCCAGAATATGACGGGCA |
| *luxS* expression reverse primer | CGCACCGGCTTTTACATGAG |
| *luxS* cloning forward primer | **CGCGGATCC**ATCGGAGGTGACTAAATGCC |
| *luxS* cloning reverse primer | **CCCAAGCTT**TGGAACCGCTTACAAATAAGAC |
| *lsrR* knockout forward primer | CAAAGTAAAGCCAGGTTATGACAATGAGCGATAATACGTTGG**CATATGAATATCCTCCTTAG** |
| *lsrR* knockout reverse primer | GAATTATTTTCCCTGCGGTTTTCTGATCGGTAACCAGTGC**GTGTAGGCTGGAGCTGCTTC** |
| *lsrR* knockout confirmation forward primer | GTGGCGTTAATCACGGTTAT |
| *lsrR* knockout confirmation reverse primer | TATTGAATTGAGGTAAGTGT |
| *lsrR* expression forward primer | GAATATCGCCTACTGCGCCT |
| *lsrR* expression reverse primer | AAATAGCGTGCGGGATGTGA |
| *lsrB* knock out forward primer | ATGGCAAGACACAGCATTAAAATGATCGCCTTACTCACTGCATATGAATATCCTCCTTAG |

| | |
|---|---|
| *lsrB* knock out reverse primer | TGAAAATGACACGCTCCGGCAATAACACAATGC CGTTACCGTGTAGGCTGGAGCTGCTTC |
| *lsrB* knockout confirmation forward primer | CGTCACCAAATCCTTGAATG |
| *lsrB* knockout confirmation reverse primer | CCACAGCCTTTTAATGCGAA |
| *lsrB* expression forward primer | AGAGTTTGGCCTGTGGGATG |
| *lsrB* expression reverse primer | GGTGAAACGGTGACTTTGCC |
| *lsrK* knock out forward primer | TGGCTCGACTCTGTACCCATACTGAATCAGGAC ATTACCTCATATGAATATCCTCCTTAG |
| *lsrK* knock out reverse primer | CGCTTTCCAGAGGGATGTCGTTAAGCCACCATC GACCAGGTGTAGGCTGGAGCTGCTTC |
| *lsrK* knockout confirmation forward primer | CGGTAACTATATCAATGCAC |
| *lsrK* knockout confirmation reverse primer | GCGGTATTCATGATTCTTCT |
| *lsrK* expression forward primer | GTGAAATGTGACCGAGCAGC |
| *lsrK* expression reverse primer | CTTTATGCTCAGCGGCGAAC |
| *cheY* expression forward primer | CTGCGGTGAACGGTTTTACG |
| *cheY* expression reverse primer | ATCTCCGACTGGAACATGCC |
| *motA* expression reverse primer | TTGCTGGCGTTGCTCTA |
| *motA* expression reverse primer | GATGATCAGGCGCAGAT |

| | |
|---|---|
| *fliM* expression forward primer | CGTCGTCAATACGCCGTTTC |
| *fliM* expression reverse primer | CACCAGGTTATCACGCCAGT |
| *fliC* expression forward primer | CTAAACAAACTGGGTGGCGC |
| *fliC* expression reverse primer | GCACCCAGGTCAGAACGTAA |
| *fljB* expression forward primer | TACGATGAAGCGACAGGAGC |
| *fljB* expression reverse primer | TTTTACCGTCTACGCCACCC |
| *fliA* expression forward primer | CGATTCGCCGACTTCCAGTA |
| *fliA* expression reverse primer | GGCGATAGCATCGAACTGGT |
| *flhD* expression forward primer | CGCCTCGGTATCAACGAAGA |
| *flhD* expression reverse primer | CACTTCATTGAGCAGACGCG |
| *hilD* expression forward primer | **CGAACCTGGGATGTTGGTGCT** |
| *hilD* expression reverse primer | **CTGGCAGGAAAGTCAGGCGT** |
| *hilA* expression forward primer | CGCCGGCGAGATTGTGAGTA |
| *hilA* expression reverse primer | TGGCGGAGACACCACTACGA |
| *invF* expression forward primer | CGTCGTTTGTGCAGCAGAGC |
| *invF* expression reverse primer | TAATTTCCGCGGCGAAACGC |

| | |
|---|---|
| *pmrD* expression reverse primer | GCAGGAATAACAGCGTGCAT |
| *pmrD* expression reverse primer | TGTTCTGGTGCTGTGCGATA |
| *pmrA* expression forward primer | AGCATCAACCGTGAGAGCAA |
| *pmrA* expression reverse primer | GTTAAACCCTTCGCCCTGGA |
| *pmrB* expression reverse primer | GGGTGCCTGTTCTTTCCGTA |
| *pmrB* expression reverse primer | TTATGGCGGTCGAAGACGAG |
| 16s rRNA forward primer | CGCTTCTCTTTGTATGCGCC |
| 16s rRNA reverse primer | TCGTCAGCTCGTGTTGTGAA |

99
